# Optoretinography reveals rapid rod photoreceptor movement upon photoisomerization

**DOI:** 10.1101/2025.03.22.644466

**Authors:** Huakun Li, Connor E. Weiss, Vimal Prabhu Pandiyan, Davide Nanni, Teng Liu, Pei Wen Kung, Bingyao Tan, Veluchamy Amutha Barathi, Leopold Schmetterer, Ramkumar Sabesan, Tong Ling

## Abstract

Rod photoreceptors are essential for vision under dim light conditions. The onset of rod-mediated vision is marked by the isomerization of rhodopsin. Here we demonstrate that human and rodent rods undergo a minute and rapid contraction of their outer segments immediately upon photoisomerization. The contraction is explained as an electro-mechanical manifestation of the rod early receptor potential generated in the disk membranes, which is challenging to access in electrophysiology. The bleach-strength dependence of the contraction was accounted by a voltage-dependent membrane tension model, developed earlier to explain a similar behavior in cones. The *in vivo* optical imaging of light-evoked electrical activity in rodent rods was facilitated by an ultrahigh-resolution point-scan optical coherence tomography (OCT) system coupled with unsupervised learning, while in humans, an adaptive optics line-scan OCT facilitated high-speed recordings in individual rods. The non-invasive *in vivo* optical imaging of rhodopsin activation will have a significant impact on diagnostics and treatment of retinal disease, especially given the vulnerability of rods in inherited and age-related macular degeneration.

## 1. Introduction

Vision begins with the photoisomerization of visual pigments^1^. Phototransduction amplification has been extensively studied for its role in the exquisite sensitivity of rods to detect single photons^2–4^. The early decline of rod function during normal aging^5^, age-related^6^ and inherited^7^ retinal degenerations has further emphasized the importance of this vulnerable cell population. Efforts to probe the functional capacity of rods include scotopic visual sensitivity^8^, rod-mediated dark adaptation^9^ and scotopic electroretinogram (ERG)^10^ *in vivo* while patch-clamp and suction-pipette electrical recordings have been used *ex vivo*^4^. Together, these assays have provided access to the various steps involved in rod phototransduction and the visual retinoid cycle^1,2,11,12^. The *in vivo* techniques are limited in their sensitivity, specificity and cellular resolution, while the *ex vivo* approaches are invasive to be used in living eyes. An ideal measure of rod viability would inform of both its structural and functional integrity with high sensitivity and resolution.

Recently, optoretinography (ORG) has emerged as a non-invasive technique to probe light-evoked photoreceptor activity, including the early steps of phototransduction, amplification and the visual cycle^13–16^. Remarkably, phase-resolved optical coherence tomography (OCT) has enabled reliable and quantitative detection of photoreceptor outer segment (OS) elongation associated with phototransduction^16–22^. Such elongation has been attributed to water influx driven by osmotic imbalance, as well as hydrolysis of the all-trans chromophore during phototransduction^16,23^. With adaptive optics (AO), these light-evoked OS responses can be localized to individual cones^18–20,24^. Interestingly, ORG has also led to the discovery that the activation of cone opsins by a strong bleaching stimulus induces a rapid (<5 ms) contraction of human cone OS preceding the later elongation^13,17,18^. The light-evoked rapid contraction of cone OS was hypothesized to result from voltage-dependent membrane tension—the electrostatic repulsions between mobile ions near the disk membrane—during the early receptor potential (ERP)^13,25^.

However, the corresponding contractile response of rod OS upon light activation has remain elusive. Previous studies have primarily focused on the light-evoked elongation of rod OS that peaks at around half a second^19,22^. Unlike cones whose disk membranes are confluent with the cell membrane and thus easily accessible for suction-pipette recordings^26^, rod disk membranes are predominantly isolated and electrically uncoupled from the plasma membrane^27–29^. Thus, conventional electrophysiology cannot readily access the ERP on the rod disk membranes, which arises from charge displacements accompanying the conformational change of visual pigments upon photoisomerization. We sought to test the feasibility of ORG as a candidate tool to access the optical manifestation of electrical activity within the rod disk membranes. Specifically, we test the hypothesis that, like cones, rod disks exhibit an ERP that produces a rapid and minute electromechanical contraction of the OS. This hypothesis was first tested in rod-dominated rat retina, starting from an unsupervised learning approach that facilitated the extraction of the light-evoked movement of the rod OS tips and the retinal pigment epithelium-Bruch’s membrane complex. Testing the hypothesis in humans entailed the use of an AO line-scan OCT instrument that was able to spatially segregate the rod and cone responses in 3-dimensions.

## 2. Results

### 2.1 Rapid movements in rod photoreceptors and adjacent outer retinal structures in response to a strong bleach

An OCT cross-sectional scan of the wild-type rat retina revealed several hyperreflective bands in the outer retina (colored bands in Fig. 1A), including the external limiting membrane (ELM), inner segment/outer segment junction (IS/OS), and a thick composite layer. The composite layer comprises the OS tips, retinal pigment epithelium (RPE), and Bruch’s membrane (BrM). Light-evoked dynamics at individual pixels in this composite layer can be extracted from the OCT phase signals using the IS/OS as a reference (see Methods). Figure 1B shows the distribution of OCT phase signals in the composite layer in response to a 34.0% bleaching stimulus, where a rapid and pronounced decrease in the OCT phase is observed, revealing two bands with distinct amplitudes (Fig. 1B). For subsequent analysis, the temporal evolution of OCT phase was converted to a change in optical path length, ΔOPL(t), using the OCT center wavelength.

**Figure 1.**
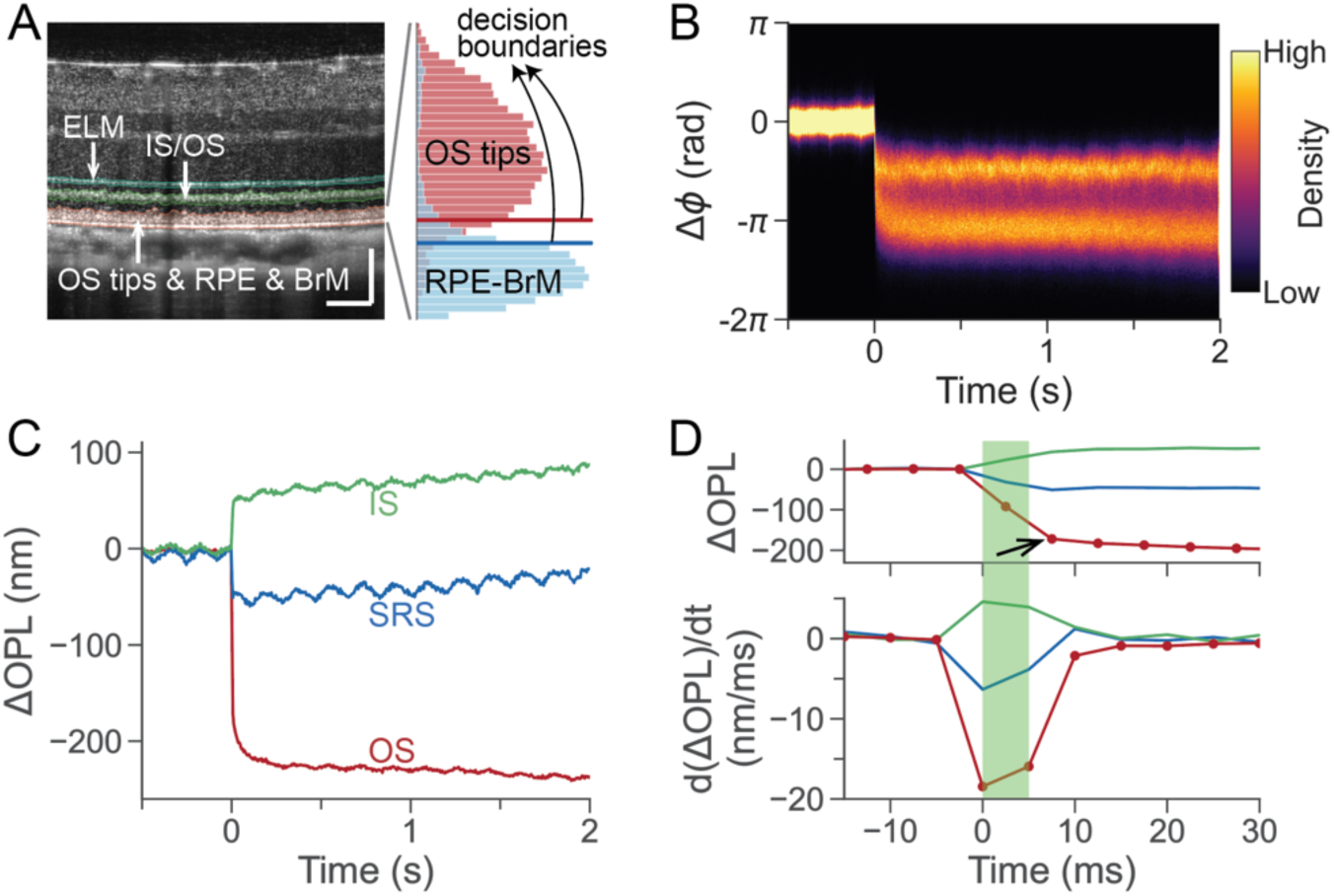
Multilayered outer retinal dynamics in response to a strong bleaching stimulus (5 ms, wavelength 505 nm, 34.0% bleach) recorded using phase-resolved OCT. (A) A representative structural image of a wild-type rat retina captured by the ultrahigh-resolution OCT. Scale bar: 100 μm. ELM: external limiting membrane; IS/OS: inner segment/outer segment junction; OS: outer segment; RPE: retinal pigment epithelium; BrM: Bruch’s membrane. Inset illustrates the isolation of OS tips and the RPE-BrM complex based on two decision boundaries (red and blue lines). (B) The distribution density of phase traces extracted from the composite layer that consists of OS tips, RPE and BrM, by taking IS/OS as the reference. The stimulus was delivered at t = 0 s. (C) Multilayered outer retinal dynamics in response to the visual stimulus. IS dynamics: ΔOPL between ELM and IS/OS; OS dynamics: ΔOPL between IS/OS and OS tips; SRS dynamics: ΔOPL between ELM and the RPE-BrM complex. (D) Top: An enlarged view of the initial rapid contractile response. Bottom: Time derivatives of the rapid responses. The green block represents the 5-ms visual stimulus.

These distinct signal patterns are categorized as arising from the OS tips and the RPE-BrM complex respectively, and are efficiently segregated using an unsupervised machine learning approach based on principal component analysis^22^ (see Supplementary Fig. 1A). However, for lower bleach levels, the signal patterns overlap, and temporal features alone are less effective in separating them (Supplementary Figs. 1B-C). To obtain a reliable criterion for clustering, we analyzed the axial distribution of the two signal types (see Supplementary Section 1 and Supplementary Fig. 2), as the anatomical structure along depth is expected to be consistent across measurements^30^. According to this criterion based on the normalized distance to the BrM, the composite layer was separated into rod OS tips, a transition region, and the RPE-BrM complex (see inset in Fig. 1A). In eight experiments with bleach levels ranging from 34.0% to 62.6%, we obtained an *F*_0.5_ score of 0.933 (0.023) [mean (standard deviation)] for OS tips using a decision boundary of 0.421 (0.027) and an *F*_0.5_ score of 0.925 (0.010) for the RPE-BrM complex using a decision boundary of 0.344 (0.026). We then used these decision boundaries to isolate OS tips from the RPE-BrM complex and calculated their respective dynamics with respect to IS/OS.

This multilayered adaptation of ORG allowed us to visualize the rapid light-evoked responses in the entire outer retina in a single measurement. As shown by the red curve in Figs. 1C-D, the ΔOPL of rod OS underwent a rapid decrease in response to the visual stimulus, while the IS dynamics, derived from the ΔOPL between ELM and IS/OS, displayed a rapid increase (green curve in Figs. 1C-D). The subretinal space (SRS) dynamics (spanning from the ELM to the RPE-BrM complex, depicted by the blue trace in Figs. 1C-D) exhibited a rapid ΔOPL decrease with a smaller amplitude compared to that of the OS. Unlike human cones, where the rapid OPL shrinkage of OS in response to a strong stimulus is typically short-lived (peaking at ∼10 ms) and of small amplitude (<50 nm) compared to the slow OS elongation^13^, the rod OS contraction in rodents was persistent over seconds and more pronounced, reaching over 200 nm (Fig. 1C) at this bleach level.

As illustrated in Fig. 2A, we further calculated the ΔOPL between the ELM and the IS/OS, OS tips, and the RPE-BrM complex immediately after the flash offset (see the time point indicated by the black arrow in Fig. 1D). We observed a contraction between the a) ELM and OS tips and b) ELM and RPE-BrM complex during the rapid OS response. Consistent with the finding in Fig. 1C, the latter was smaller in magnitude. Surprisingly, we also found a rapid increase in the ΔOPL along the IS (from ELM to IS/OS) during this period. A plausible interpretation for these changes in the outer retina is that the OS undergoes an intrinsic mechanical shrinkage that reduces the ΔOPL in the OS; such a movement would pull the IS posteriorly via the IS/OS junction and the RPE-BrM complex anteriorly via connected microvilli^31,32^. This interpretation is consistent with the smaller magnitude of contraction in the SRS (ELM – RPE/BrM) compared to that between the ELM and OS tips. An alternate interpretation may be a contribution from refractive index changes, which is considered further in the Discussion.

**Figure 2.**
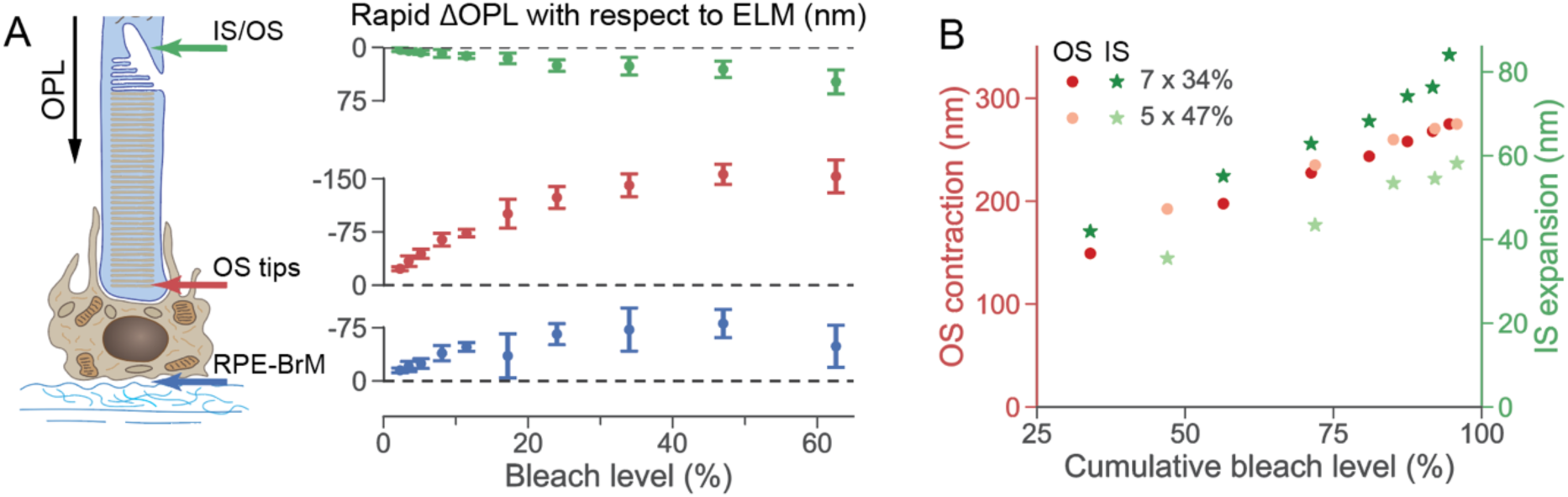
Rapid ΔOPL in the major outer retinal layers at varying bleach levels. (A) ΔOPL measured from the ELM to the IS/OS (green), OS tips (red), and the RPE-BrM complex (blue) immediately after the 5-ms flash offset. Data points and error bars represent the mean and standard deviation respectively at a specific bleach level. (B) The rapid movement of rod IS (green stars) and OS (red dots) in response to serial flashes (5 or 7 flashes, each yielding a 47% or 34% bleach level per flash of the remaining photopigment). The flash duration and inter-flash interval were 5 ms each. Data points (5 or 7) represent the successive OS & IS movement under the two conditions. OPL: optical path length; ELM: external limiting membrane; IS/OS: inner segment/outer segment junction; IS: inner segment; OS: outer segment; RPE: retinal pigment epithelium; BrM: Bruch’s membrane.

We conducted multilayered ORG recordings under serial-flash stimuli to test the correspondence between the IS expansion and OS contraction. Figure 2B shows the rod IS and OS movements after each series of either 5 or 7 flashes (see Supplementary Fig. 3). In both cases, each flash lasted 5 ms with an interval of 5 ms between the adjacent flashes. In total, one set of serial flashes lasted 45 ms (5 flashes + 4 intervals), and the other lasted 65 ms (7 flashes + 6 intervals), while both aimed at reaching the same cumulative bleach level (∼95% bleach). Under the serial-flash stimuli, the IS expansion was correlated in its bleach dependence with the OS contraction, suggesting that these two rapid responses are initiated by the same underlying process. While both serial-flash stimuli triggered nearly identical rapid OS contractions resulting from the same cumulative bleach level, variability was observed in the amplitude of the rapid IS expansion between these two measurements. This difference could be attributed to the larger inter-individual variation observed in the IS expansion compared to the OS contraction. For instance, measurements in seven rats showed that in response to a single flash at a bleach level of 62.6%, the rapid OS contraction had a standard deviation of 13.6 nm (mean value: 201.8 nm), while the rapid IS expansion had a standard deviation of 16.9 nm (mean value: 48.5 nm). Thus, the smaller magnitude, more variable IS expansion may not be an ideal substitute for the OS contraction.

#### A voltage-dependent membrane tension model explains the rapid contraction in rod OS accompanying rhodopsin activation

In previous studies on the rapid light-evoked contraction of human cone OS, a logarithmic relationship between the contraction amplitude and the stimulus strength was reported^13,25^, which is in line with the prediction of a voltage-dependent membrane tension model^25^. We investigated whether a similar model is able to explain the bleach dependence of the rapid rod OS contraction.

Figure 3A shows representative dynamics of the rod OS evoked by visual stimuli at bleach levels ranging from 2.3% to 62.6%. It is important to note that the lowest bleach level used in this experiment (>2 %) was still considerably higher than those used in our previous study on light-evoked rod OS elongation (<0.3% bleach level)^22^. In response to the strong bleaching stimuli, a rapid contraction was observed during the 5-ms flash stimuli (enlarged view in Fig. 3A), with the amplitude increasing with the bleach level. An elbow was consistently observed at 7.5 ms post-stimulus (the averaged timestamp of the B-scan after the flash offset, indicated by the blue arrow in Fig. 3A), where the first, rapid contractile kinetics gives way to a slower response. For bleach levels between 2.3% and 11.5%, the rapid contraction in the OS was followed by a continuous, slow elongation that persisted over the 37-second recording. Under stronger stimuli (>11.5% bleach), we observed a secondary contraction with slower kinetics than the rapid contraction in the first 7.5 ms, peaking at 5 ∼ 15 s. This secondary contraction was followed by continuous elongation, similar to the signals observed under 2.3% - 11.5% bleach levels. For clarity in this article, we categorized these processes as rapid OS contraction (within 7.5 ms), secondary OS contraction (if any, from 7.5 ms to ∼10 s), and late OS elongation.

**Figure 3.**
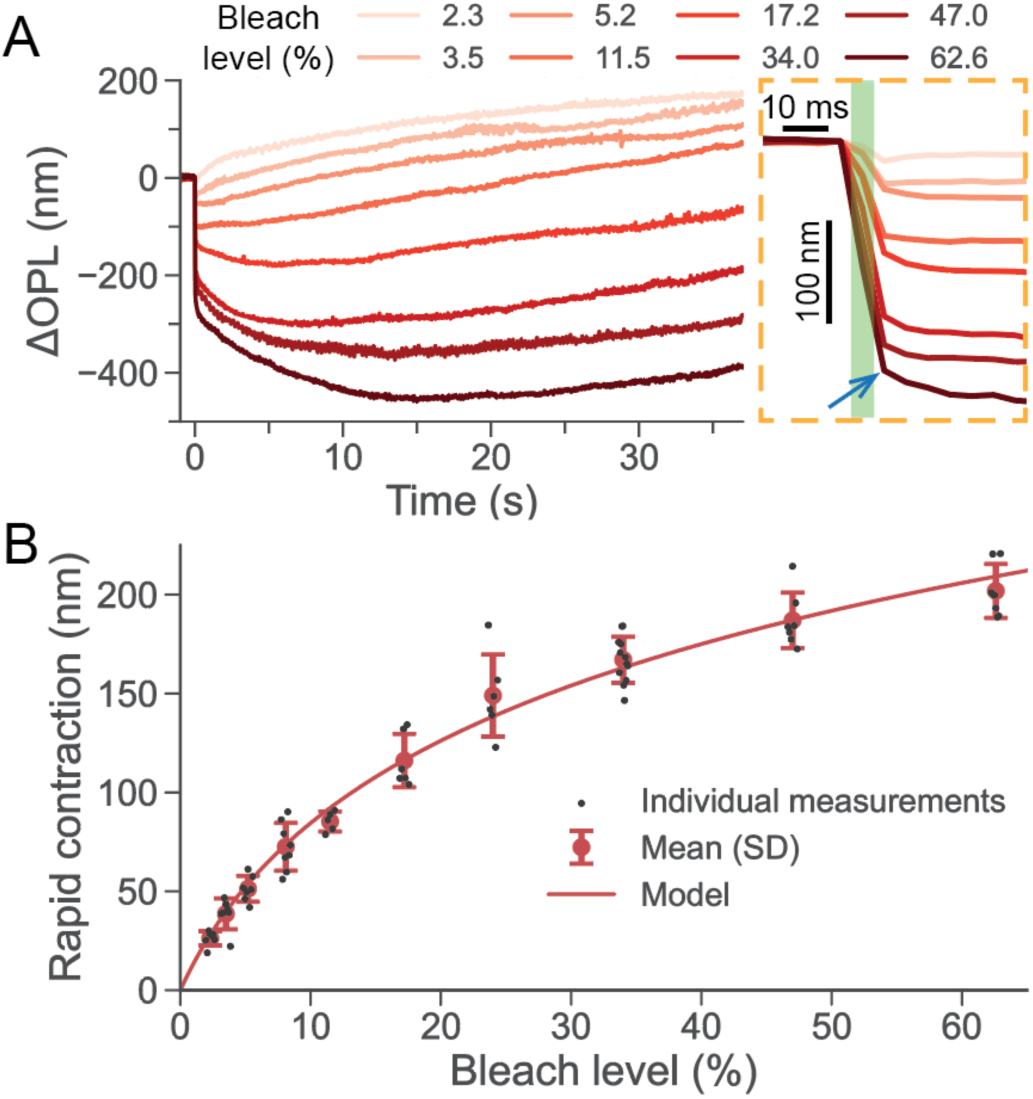
Dependence of the light-evoked rod OS dynamics on the stimulus strength. (A) Representative traces of the OS dynamics in response to flashes ranging from 2.3% to 62.6% bleach levels, with the enlarged view on the right showing an elbow point at t = 7.5 ms after the flash onset. (B) The rapid OS contraction amplitude at varying bleach levels. Black dots represent individual measurements. The red circles and error bars denote the mean values and corresponding standard deviation (SD) ranges at each bleach level, respectively. The red curve represents the prediction from the voltage-dependent membrane tension model.

Although the exact onset of the rapid OS contraction could not be measured due to the limitation in our imaging speed, its millisecond time scale coincides with that of rhodopsin activation, transitioning from rhodopsin to metarhodopsin II (Meta II)^1^. For the rapid OS contraction to be induced by osmolarity-driven water flow, the water permeability coefficient of the cell membrane would need to be 0.30 cm⁄s (see Supplementary Section 2), which is much larger than the 2.6 × 10^$%^ cm⁄s measured in in-vitro studies^33^. Thus, the observed rapid OS contraction is too fast to be induced by osmosis.

Since the rhodopsin activation is accompanied by charge displacement across disk membranes, i.e. the ERP^28,34–37^, we tested whether electromechanical coupling could be responsible for driving the rapid rod OS contraction. A voltage-dependent membrane tension model was previously proposed to explain the rapid contraction in cone OS^25^ as well as cellular deformations in mammalian neurons during their millisecond-scale action potentials^38–40^. In photoreceptor OS, the hyperpolarization of disk membranes during the R2 phase of the ERP leads to an increase in the repulsive forces between mobile ions within the Debye layers^35,41^, causing the disk membrane to stretch. Under physiological conditions, the normalized area expansion of the disk membrane (Δ*A*⁄*A*_0_) at different membrane tensions can be calculated by^25,42^,

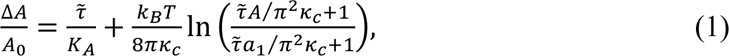

where *A*_0_ is area of the disk membrane patch when the observable tension *τ̃* is 0, *A* and Δ*A* are tension-dependent membrane area and the corresponding area expansion, respectively, *k_B_* is the Boltzmann’s constant, *T* is the absolute temperature (310 K). *k_A_* is the area expansion modulus, *K_A_* is the bending modulus, and *k_c_* is the smallest bending feature size.

Assuming volume conservation within the millisecond-scale dynamics, the lateral expansion of individual disks would result in their axial contraction. The contraction of the rod OS during the ERP was thus modeled as the cumulative contraction of individual disks, as detailed in Supplementary Section 3. Interestingly, by fitting the parameters listed in Table 1, the modeled rapid contraction amplitudes matched the experimental observations across varying bleach levels (see the red curve in Fig. 3B). Furthermore, all the fitted parameters (*τ̃*_0_, *k_c_* and *a*_1_) fall into the typical ranges from literature. Overall, the voltage-dependent membrane tension model can explain the observed rapid rod OS contraction as being caused by the ERP in the disk membranes.

**Table 1.** Fitting parameters for the voltage-dependent membrane tension model.

| Parameter | Fitted value | Typical value<sup>25</sup> | Unit |
| --- | --- | --- | --- |
| Initial tension, $\tilde{\tau}_0$ | 0.13 | 0.1 – 1 | $\mu\text{N/m}$ |
| Bending modulus, $\kappa_c$ | $0.38 \times 10^{-19}$ | $0.5 - 2 \times 10^{-19}$ | $\text{N}\cdot\text{m}$ |
| Smallest patch size, $a_1$ | 23 | 25 | nm |

### 2.2 Rod OS responses elicited by multi-flash stimulation

Based on the hypothesis linking the rod OS contraction to the ERP within the disk membranes, it is predicted that a single flash or a set of serial flashes will both elicit the same amplitude of contraction, provided they confer the same effective rhodopsin bleach. Further, under both stimulation conditions – single or serial flashes – the saturation of the contraction amplitude should be explained by the same voltage-dependent membrane tension model. We tested this prediction by undertaking prolonged ORG with multiple flashes of equivalent strength that were separated by a longer time interval compared to Fig. 2B. Figure 4A illustrates the rod OS responses to 5 flashes with an inter-flash interval of 15 s, where each flash bleached 5.2% of the remaining rhodopsins. To better compare the rod OS responses evoked by consecutive flashes, we aligned the signals extracted from each epoch to their flash onsets (see Fig. 4B). Notably, we observed that successive flashes significantly slowed the rate of late OS elongation. The slowdown is particularly evident after the 1^st^ flash, where only a small fraction of rhodopsin (5.2%) was bleached. Supplementary Fig. 4 shows another similar example with each flash bleaching 8.1% rhodopsins. This observation suggests that the late OS elongation, purported to be linked to the amplification stages of the phototransduction cascade^16,23^, is heavily suppressed by the first flash. This phenomenon resembled previous observations using electrical recordings, where a second flash, delivered 1 min after the first flash, could isolate the ERP from the late receptor potential^35,37^. It follows that the enzymes and/or substrates underlying the phototransduction cascade and the late OS elongation are temporarily exhausted by the first flash, such that the later flashes are unable to activate the amplification cascade and the resultant OS elongation with the same velocity.

**Figure 4.**
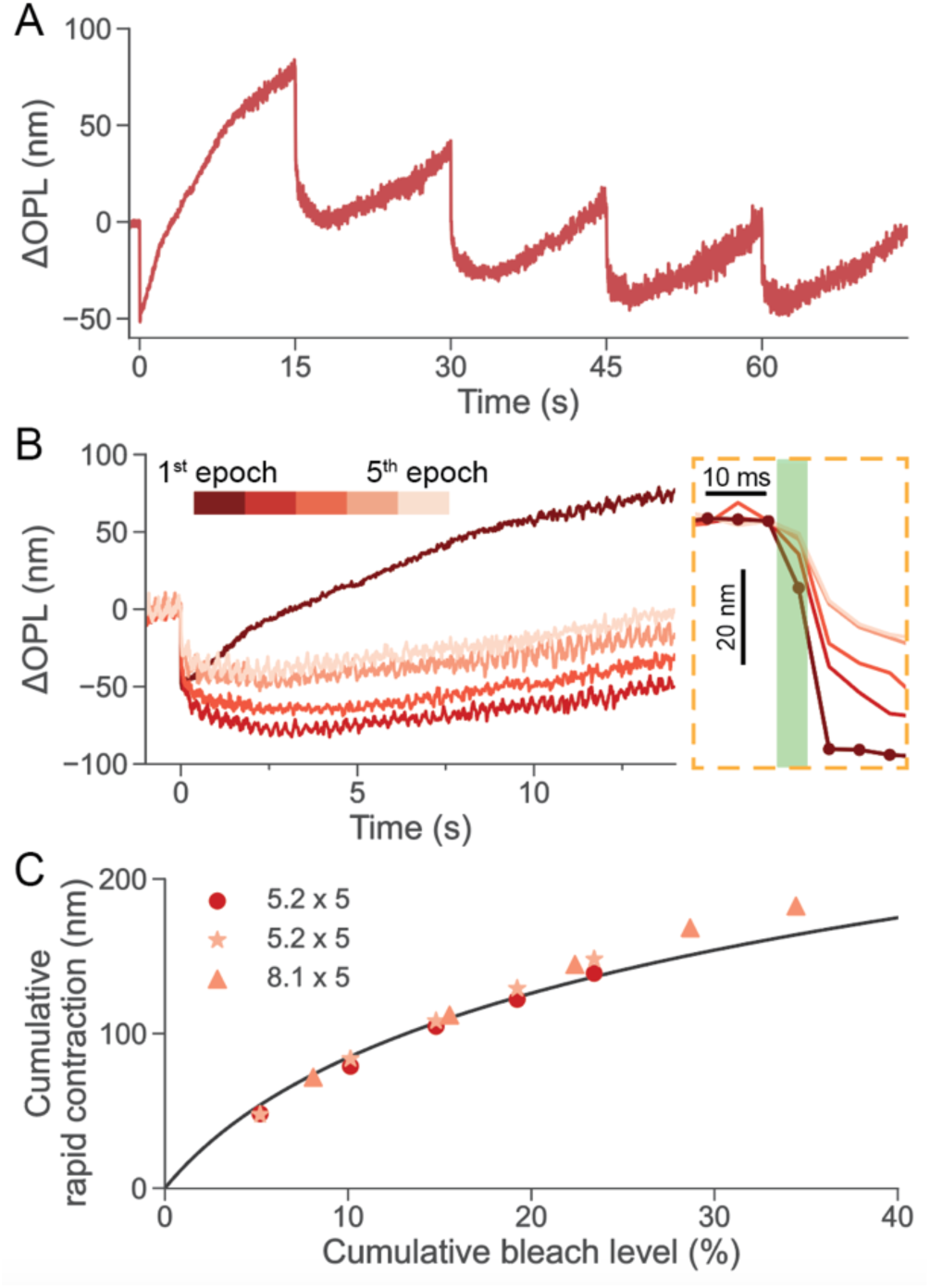
Rod OS responses evoked by 5 flashes with an inter-flash interval of 15 s. (A) Prolonged ORG recording of the rod OS responses to five flashes over 75 s, with each flash bleaching 5.2% of the remaining rhodopsins. (B) The rod OS responses extracted from each epoch were aligned to the flash onsets, with an enlarged view of the rapid OS responses shown in the dashed rectangle. (C) The cumulative rapid contraction, calculated as the sum of the rapid OS response amplitudes elicited by all flashes, was plotted against the cumulated bleach level, which represents the total percentage of rhodopsins bleached by the consecutive flashes. Dots correspond to the measurement in Fig. 4A-B, and star markers were extracted from the measurement conducted in another animal following the same protocol. Triangle markers correspond to the measurement in Supplementary Fig. 4. The black curve represents the same modeled curve in Fig. 3B.

On the other hand, the quantitative relationship between the rapid contraction and the bleach level remained unaffected by the alteration of the late OS elongation properties after the first flash. We calculated the cumulative rapid contraction by summing the rapid contraction evoked by all flashes (Fig. 4C). As predicted by the hypothesis, the relationship between the cumulative rapid contraction and the corresponding bleach levels aligned well with the voltage-dependent membrane tension model (the black curve in Fig. 4C, identical to the modeled curve in Fig. 3B).

### 2.3 Rapid contraction in human rod OS in response to strong bleaching stimuli

Next, we conducted ORG experiments in human subjects using an AO line-scan OCT system^43^ to explore whether the rapid rod OS contraction observed in rodents also occurs in humans. Figure 5 illustrates ORG signals measured from the 10° temporal retina at a volume rate of 41.4 Hz. In the en-face image at the rod OS tips (Fig. 5A), individual rod photoreceptors can be identified (see the enlarged view in Fig. 5B). In response to a 5-ms, 470 nm stimulus that bleached 27.1% of rhodopsin, we observed consistent rapid contraction (t = 10 ms) and late elongation (t = 1.7-1.9 s) in individual rod OS (colored dots in Fig. 5B). Figures 5C and 5D shows the rod OS responses evoked by visual stimuli of varying strengths. At a 0.6% bleach level, the rod OS exhibited a typical elongation response, consistent with the previous reports^19,22^. Interestingly, at higher bleach levels, we observed a rapid contraction preceding the late elongation. To enhance the temporal resolution, we divided each volume into five sub-volumes along the slow scan dimension and extracted individual rod ORG signals from each sub-volume. As shown in Fig. 5E, the contraction amplitude gradually increased with bleach strength, with peak contraction occurring 2-6 ms post-stimulus. The contraction amplitude saturated at ∼100 nm for bleach levels above 27.1%. On the other hand, when inspected even with the increased temporal resolution (200 Hz), the lowest bleach strength tested (0.6 %) did not show an observable contraction.

**Figure 5.**
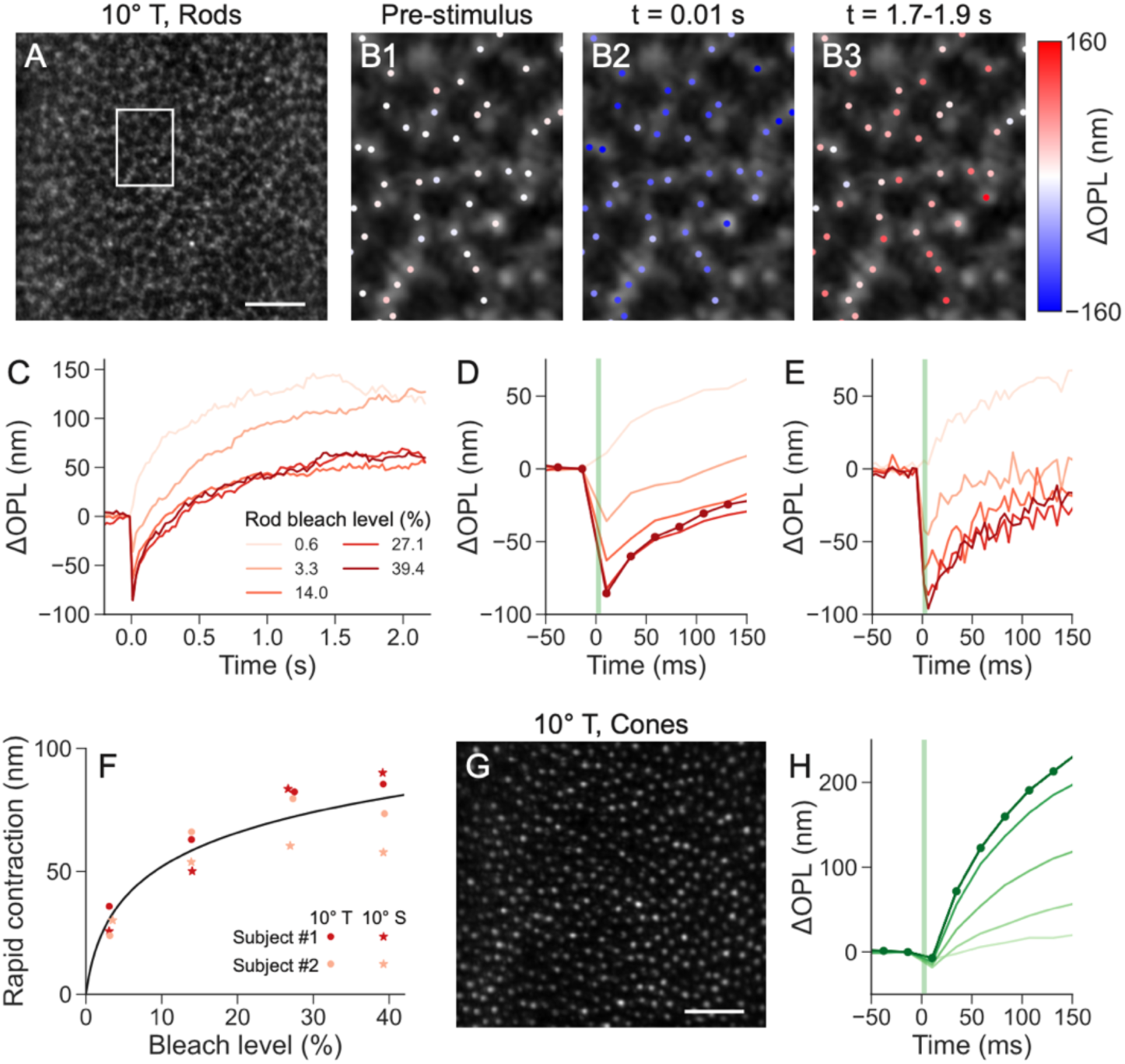
ORG responses in human rod and cone OS evoked by visual stimuli at varying strengths. (A) Maximum intensity projection at rod OS tips at 10° temporal eccentricity. Scale bar: 50 µm. (B) Enlarged view of the rod OS tips (enclosed white region in Fig. 5A). The colored dots denote manually selected individual rods, with color grading indicating the ΔOPL along the rod OS. (B1) ΔOPL extracted from the volume right before the visual stimulus, (B2) ΔOPL extracted from the volume immediately post the stimulus, indicating rapid contraction, and (B3) ΔOPL averaged over a period 1.7-1.9 s after the visual stimulus, indicating late elongation. (C) Rod OS signals in response to stimuli bleaching 0.6%-39.4% rhodopsins. (D) The enlarged view of the rod OS contraction. (E) Corresponding rod OS responses extracted using an ultrafast analysis method, where temporally sub-sampling each volume into five sub-volumes along the slow scan dimension increases the sampling rate to ∼200 Hz. (F) Amplitude of rapid rod OS contraction showed a logarithmic dependence on the bleach level (black curve). The individual data points were extracted from ORG signals shown in Supplementary Fig. 5. Small jitters were introduced to the horizontal axis to reduce overlap between data points. (G) Maximum intensity projection at cone OS tips from the same region as in Fig. 5A. Scale bar: 50 µm. (H) Cone OS responses extracted from the same experiment. Stimulus strengths ranged from 2.8 × 10^’^ − 2.7 × 10^(^ photons⁄µm^2^, corresponding to 0.6% (light green) to 39.4% (deep green) rhodopsin bleach (see legend in Fig. 5C and Supplementary Table 4).

The AO-OCT 3D volume acquisition enables sectioning at different depths to selectively reveal the cone or the rod mosaic structure and corresponding ORG for the same visual stimulus. Figures 5G and 5H display the en-face image at the cone OS tips and the associated ORG signals extracted from the cone OS for comparison to the rods. The contraction in the cone OS is significantly smaller, while the elongation is substantially larger and more rapid in its rate of increase, in comparison to the rod OS.

Figure 5F shows that the amplitude of rapid rod OS contraction increased logarithmically with increasing bleach levels, similar to measurements in rodent rod OS. This suggests that the rapid rod OS contraction observed in rodents and human subjects shares similar underlying mechanisms. However, there exist obvious qualitative differences between species. First, the amplitude of rod OS contraction is substantially greater in rodents for the same bleach strength. Second, the recovery of the OS contraction is significantly faster in human subjects. This faster recovery indicates that a time-dependent model may be required to describe the bleach-strength dependence of contraction amplitude measured in human rod OS^25^.

## 3. Discussion

Ex vivo studies in rods have reported rapid structural changes and optical signatures upon exposure to visual stimuli. For instance, studies in intact frog retina and individual osmotically intact rod OS have revealed a birefringence loss occurring at a rate aligned with Meta II formation^44,45^. In addition, light scattering experiments in suspensions of fragmented bovine rod OS have reported signals corresponding to the intermediate steps of phototransduction^46^. However, the models proposed to explain such changes have been inconsistent with each other^44,45,47,48^. Here, we discovered that rodent and human rod OS undergo a rapid OS contraction upon photoisomerization whose magnitude scales logarithmically with bleach strength (Fig. 3 & 5). Additionally, we observed that the OS contraction was synchronous with an expansion of the rod IS (Fig. 2 & Supplementary Fig. 3). Using serial flashes, we further confirmed that the rod OS contraction and elongation have fundamentally different kinetics and light sensitivity, such that a first priming flash can readily segregate the two components (Fig. 4). Finally, a comparison of cone and rod responses in the same human subjects and retinal locations undertaken with AO revealed substantial differences in kinetics, light sensitivity and magnitude of OS length change (contraction and expansion) between species and photoreceptor types.

The observed rapid OPL decrease in the OS may stem from changes in the refractive index, the physical length, or a combination of both. Thus, it is important to delineate the relative contributions of both factors in the observed OS contraction. With respect to refractive index changes, it can arise in the rod OS 1) from a loss in visual pigment-related anomalous dispersion following rhodopsin activation. However, this effect was found to be negligible for wavelengths longer than 650 nm^45,49^, which covers the wavelengths used in our OCT imaging (750 nm-950 nm). 2) Refractive index changes may also occur during the reorientation of anisotropic molecules within the disk membranes and alternations in the arrangement of lamellar dielectric structures that comprise disk membranes, cytoplasmic and intradisk spaces^44,45^. The former would result in intrinsic birefringence changes, whereas the latter can lead to variations in form birefringence. As a reference for their scales, rhodopsin activation induced a decrease in birefringence of 7 × 10^$#^ in osmotically intact frog rod OS at wavelengths longer than 650 nm^45^, which can only account for an OPL change of a few nanometers, substantially smaller than the > 100s of nm rod OS contraction shown here. Importantly, we observed a OPL increase in the IS as well as that between the rod OS and the RPE-BrM complex (Fig. 2), which cannot be explained solely by the refractive index changes within the OS. Our results are consistent with the notion that mechanical movement is the main contributor to the observed, rapid OS length contraction accompanying rhodopsin activation.

While this study observed a similar rapid contraction in the rod OS in both rodents and human subjects, there were notable differences in the characteristics of the contraction between the two species. In rodents, the contraction was significantly more persistent, and took longer to recover compared to the human rod OS. Furthermore, the secondary contraction of the rod OS, which was prominent in rodents at higher bleach levels (>15%), was largely absent in the human rod OS response. A few previous studies have touched on the possibility that a substantially decreased dark current could result in an osmotic imbalance that shrinks the rod OS^50,51^. However, the precise mechanisms underlying the secondary contraction in rodents and the observed discrepancies between rodent and human rod OS responses remain to be elucidated. To better understand these differences, future studies may investigate the potential influence of dark adaptation protocols (12 hours in rodents versus 15 minutes in human subjects) on the rod OS response. Broadly, comparative studies exploring the structural and physiological differences in rods among various species may provide valuable insights into the factors contributing to these differences.

In human subjects, the amplitude of rapid contraction in both rod and cone OS saturated logarithmically with increasing bleach levels^13^. However the maximum contraction amplitude measured in rod OS reached up to ∼100 nm, which is significantly larger than the ∼50 nm measured in cone OS^13^. One notable difference between primate cone and rod photoreceptors is that the length of rod OS is significantly greater than that of cone OS^52,53^. The accumulation of disk membrane contraction will thus be greater along a longer OS in rods compared to cones. Additionally, the duration of the contraction period in human rod OS persisted much longer than that observed in human cone OS (hundreds of milliseconds compared to less than 20 milliseconds)^13^. This could be attributed to a roughly 5-fold lower initial velocity of human rod OS elongation, as determined by the ultrafast analysis (maximum slope = 0.8 nm⁄ms), that delays the truncation of the OS contraction. In contrast, the maximum initial cone OS elongation velocity is noted to be significantly faster (∼4 nm⁄ms)^13,16^.

Although ERP signal has been recorded from both rods and cones^28^, it is important to note that in rods, only a small fraction of rhodopsins (estimated to be 2% of the total rhodopsins in mouse rods^29^) present in the plasma membrane and the small number of nascent, basal disks can contribute to the measured electrical signal^28^. As such, despite the abundance of rhodopsin in the human and monkey retina, the ERP measured in these species using electrical approaches were primarily from the activation of cone opsins^54,55^. Remarkably, the rapid rod OS contraction reported in this study is likely to reflect the cumulative contraction of individual sealed disks, which are electrically uncoupled from the plasma membrane and thus difficult to access in electrophysiology^1,27^. The measured bleach-strength dependence of the contraction aligned with the model prediction based on the voltage-dependent membrane tension during ERP, whether assessed by a single flash (Fig. 3B) or a serial bleach protocol (Fig. 4C). The latter has been widely used in studies of the ERP, where an exponential decay in the electrical potential was observed due to a similar exponential reduction in the rhodopsin concentration following preceding flashes^28,29,36^. Unlike the linear relationship noted between the ERP and bleached pigment in prior studies^35,36^, the rod OS contraction exhibits a logarithmic saturation as described well by the voltage-dependent membrane tension accompanying electrical activity.

Overall, label-free optical imaging of the rapid rod OS dynamics in vivo can uncover the first steps in rhodopsin activation that was previously challenging to access even in electrical recordings. Importantly, the objective, non-invasive and spatially-localized monitoring of rod photoreceptor function will enable the sensitive assessment of impaired dark adaptation and visual retinoid cycle in retinal diseases.

## 4. Methods

### 4.1 Ultrahigh-resolution point-scan OCT system for rodent retina imaging

The ORG experiments were performed using a custom-built spectral-domain ultrahigh-resolution OCT system (Fig. 6A)^22^. The system employed a superluminescent diode (cBLMD-T-850-HP-I, Superlum, Ireland) with a center wavelength of 840 nm and a full-width at half-maximum (FWHM) of 146 nm, providing a theoretical axial resolution of 2.0 µm in tissue. A spectrometer equipped with a line-scan camera (CS800-800/300-250-OC2K-CL, Wasatch Photonics, USA) captured spectral interferograms at a line-scan rate of 250 kHz and allowed an imaging depth of 1.1 mm in air. An afocal telescope, consisting of a scan lens (80 mm doublet) and an ocular lens (30 mm + 25 mm doublets), conjugated the galvo scanner and the pupil plane with a magnification of 0.17. This arrangement reduced the OCT beam size and enlarged the scan angle. Based on a standard rat eye model^56^, the theoretical diffraction-limited beam size on the retina was estimated to be 7.2 µm (FWHM).

**Figure 6.**
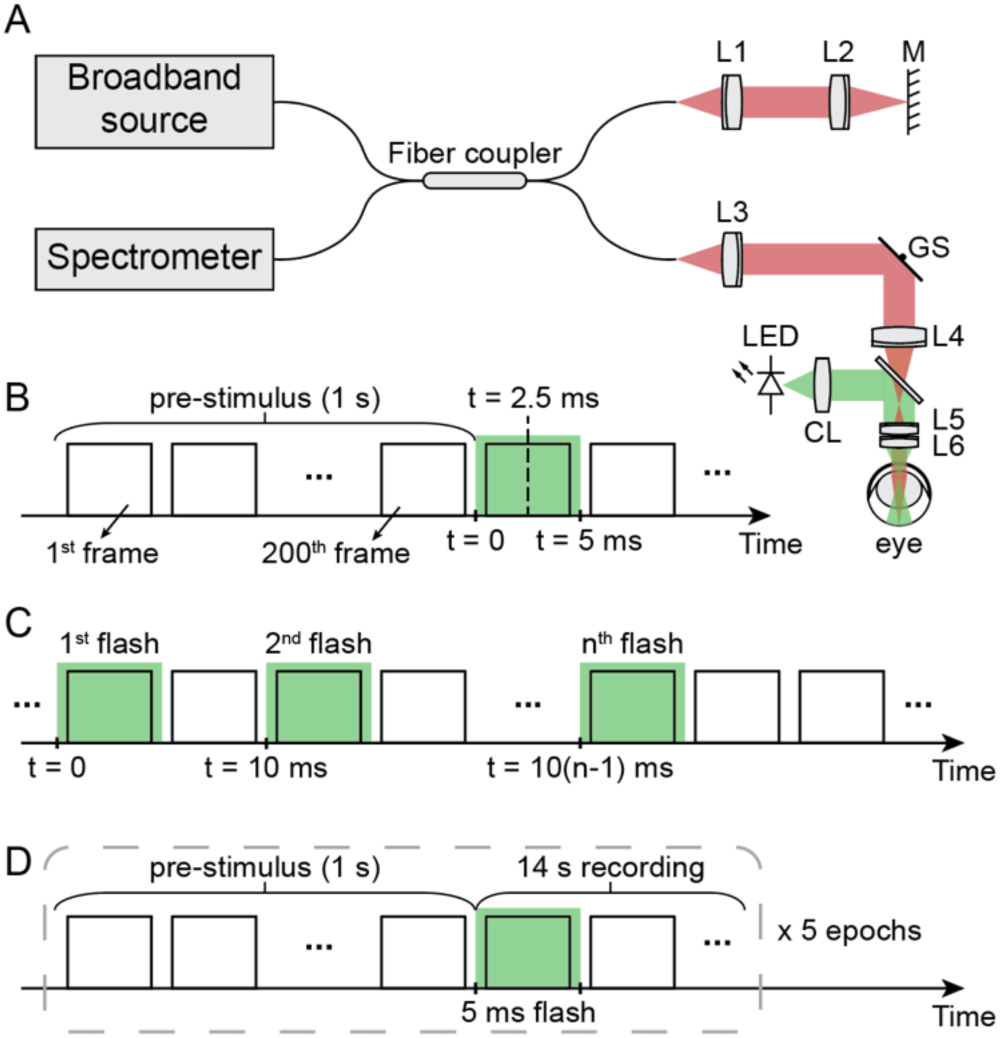
System setup and the timing diagrams for data acquisition and visual stimulation. (A) A spectral-domain point-scan OCT system (red path) was integrated with a visual stimulation channel (green path). L1-L6: doublet lenses. CL: condenser lens. GS: galvo scanner. M: mirror. (B) Protocol I: a 5-ms flash was delivered 1 s after the start of the OCT recording. (C) Protocol II: multiple 5-ms flashes were delivered at time points of 0, 10 ms, …, 10(n-1) ms, where n is the total number of flashes. (D) Protocol III: five epochs, each consisting of a 15-s recording, were captured consecutively. A 5-ms visual stimulus was delivered 1 s after the start of each epoch.

A green LED (M505L4, Thorlabs, USA) was collimated by an aspheric condenser lens to provide the visual stimulation. The ocular lens converged the collimated beam and achieved a 5.82 mm^6^Maxwellian illumination on the posterior eye. The visual stimulus settings were converted into bleach levels based on the previously published method^57,58^, as detailed in Supplementary Section 4.1. A function generator (PCIe-6363, National Instruments, USA) synchronized the camera acquisition, galvanometer scanning, and visual stimulation. Data acquisition was controlled using an interface software developed in NI LabVIEW (19.0.1, National Instruments, USA).

### 4.2 Animal preparation and ORG imaging

All experiments were conducted in compliance with guidelines and approvals from the Institutional Animal Care and Use Committee (IACUC), SingHealth (2020/SHS/1574). Wild-type Brown Norway rats (N = 13) were used in the study, and all imaging sessions were performed when they were between 7 and 17 weeks of age. The animals were dark-adapted overnight before each ORG experiment, and the entire experiment was conducted in a darkroom. During the experiment, animals were anesthetized with a ketamine and xylazine cocktail and secured in a custom-built stereotaxic stage to minimize eye motion. Before imaging, one drop of 1% Tropicamide (Alcon, Geneva, Switzerland) and one drop of 2.5% Phenylephrine (Alcon, Geneva, Switzerland) were applied to the eye to dilate the pupil. The cornea was kept moist during the imaging by frequently applying a balanced salt solution.

Repeated cross-sectional scans, each consisting of 1000 A-lines acquired in 4 ms, were collected from the retina. We investigated ORG signals evoked by a single flash at varying bleach levels (2.3% - 62.6%, Protocol I), and ORG signals in response to serial flashes with (Protocol II) short inter-flash intervals and (Protocol III) long inter-flash intervals. Note that to avoid interference from preceding bleaching, each eye was used for only one acquisition per imaging session.

In Protocol I, a 5-ms visual stimulus was delivered to the retina 1 s after the start of the OCT recording (see Fig. 6B). To capture the rapid dynamics evoked by the flash, the first 600 B-scans were acquired at a temporal resolution of 5 ms (4 ms acquisition + 1 ms galvo flyback time, i.e., a duty cycle of 0.8) for 3 s. To record the prolonged dynamics while reducing computational load, we continued the recording with 1400 B-scans at a temporal resolution of 25 ms (4 ms acquisition + 21 ms galvo flyback time, i.e., a duty cycle of 0.16) for 35 s. The total recording time was 38 s, including a 1-s pre-stimulus period. In all experiments, the onset of the first flash was set as time point 0.

Protocol II used the same OCT recording configuration as Protocol I, but instead of a single flash, multiple 5-ms flashes were delivered 1 s after the start of the OCT acquisition (see Fig. 6C). The flashes were delivered at time points of 0, 10 ms, …, 10(n-1) ms, where n is the total number of flashes.

In Protocol III, the entire recording consisted of five consecutive epochs (see Fig. 6D). Each epoch comprised 600 B-scans at a higher temporal resolution (5 ms) for 3 s and 480 B-scans at a lower temporal resolution (25 ms) for 12 s. A 5-ms visual stimulus was delivered 1 s after the start of each epoch.

### 4.3 Preprocessing of OCT images and extraction of phase traces for ORG measurements in rodents

The raw spectral interferograms first underwent spectral calibration and k-linearity. Then, a discrete Fourier transform (DFT) was applied to generate complex-valued OCT images. To minimize signal decorrelation caused by bulk tissue motion, we used our proposed phase-restoring subpixel image registration method to register complex-valued OCT images with subpixel precision^59^. Specifically, the subpixel-level translational displacements between subsequent frames and the first frame were determined using the single-step DFT algorithm^60^. Then, the axial and lateral displacements were corrected by multiplying exponential terms in the spectral and spatial frequency domain, respectively.

To detect light-evoked retinal dynamics, we calculated the temporal phase change between two retinal layers. First, we removed the arbitrary phase offset by calculating the phase change with respect to the first B-scan,

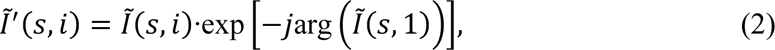

where *Ĩ*(*s*, *i*) represents the complex-valued OCT signal after image registration. *s* and *i* denote the spatial and temporal indices, respectively. *j* is the imaginary unit. arg() denotes the calculation of the argument. *Ĩ*′(*s*, *i*) is the corresponding time-referenced signal.

We further performed self-referenced measurements to remove residual bulk phase error, resulting in phase differences between two retinal layers. Specifically, for each pixel of interest in the target layer, referred to as *s_t_*, its temporal phase change was calculated relative to a reference region chosen from a reference layer. The reference region was centered on the same A-line as the pixel of interest and spanned across 5 adjacent A-lines. The phase change of the reference region, *φ^r^*, was obtained by spatially averaging the time referenced OCT signals and extracting the phase component,

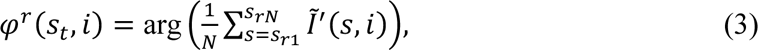

where *s_r_*_1_, *s_r_*_2_, …, *s_rN_* represent the spatial index of individual pixels in the selected reference region, and *N* is the total number of pixels in the reference region.

By referring to the reference region, the phase change of the pixel *s*_8_, denoted as *φ_tar_*_/*ref*_(*s_t_*, *i*), can be calculated as follows,

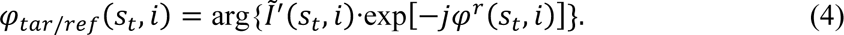

We applied the same process to every pixel in the target layer to obtain spatially resolved phase traces. Traces with a pre-stimulus standard deviation less than 0.5 radians were retained for further analysis. Note that to enhance the signal-to-noise ratio (SNR), phase traces extracted from individual B-scans spanning over 4 ms were spatially averaged within each signal pattern based on the Knox-Thompson algorithm^61^. We assigned the midpoint of the 4-ms recording as the timestamp for the averaged phase traces extracted from the corresponding B-scan. For example, the first B-scan recorded after the flash onset at 0 ms spanned from 0.5 ms to 4.5 ms, and the corresponding averaged phase traces were assigned a timestamp of 2.5 ms (see Fig. 1B). The obtained phase change can be further converted into ΔOPL by multiplying it by *λ_c_*⁄4*π*, where *λ_c_* is the center wavelength of the OCT system.

### 4.4 ORG experiments in human subjects using an AO line-scan OCT system

Two male subjects free of retinal disease were enrolled in this study. All subjects provided written informed consent before participating. An AO line-scan OCT system, as described in a previous study^16,43^, was used for human ORG experiments. The experimental protocol was approved by the University of Washington Institutional Review Board, and all procedures were in accordance with the tenets of the Declaration of Helsinki.

Cycloplegia was introduced using tropicamide 1% ophthalmic solution (Akorn Inc.) before OCT imaging to dilate the pupil. The subjects underwent a 15-minute dark adaptation before each imaging session that replenished up to 90% of rhodopsin^11^. Each session consisted of two recordings: one centered at 10° temporal and the other at 10° superior. The visual stimulation field was sufficiently small to ensure no overlap between the two regions. Two experimental protocols were employed:

*Protocol 1*: 100 volumes, each consisting of 500 B-scans and covering a 0.8°×1° field of view (FOV) were recorded at a volumetric rate of 41.4 Hz.

*Protocol 2*: 50 volumes, each consisting of 600 B-scans and covering a FOV of 0.92°×1°, were recorded at a volumetric rate of 17.3 Hz.

In both recording protocols, a 5-ms visual stimulus generated by a 470 nm LED was delivered in Maxwellian view at the start of the 11th volumetric scan. Stimulus parameters were converted into bleach levels, as detailed in the Supplementary Section 4.2. Each visual stimulus condition was repeated five times to enhance the SNR of the measured ORG.

### 4.5 Extraction of ORG signals in human ORG measurements

OCT data were processed using conventional techniques as described previously^16,43^. The acquired interference fringe data were resampled in k-space and Fourier-transformed along the wavenumber dimension, resulting in complex-valued OCT volumes. Each retinal volume was segmented, and en-face images were generated by taking maximum-intensity projections centered at the cone OS tips. En-face images were registered using a strip-based registration algorithm^62^, with lateral shifts from registration and axial shifts from segmentation being applied to register all volumes. Individual cones and rods were manually selected based on the maximum intensity projections of the cone OS tips and rod OS tips, respectively.

To measure ORG signals, we first referenced registered OCT volumes to the mean of all pre-stimulus volumes to remove arbitrary phase offsets. Next, we multiplied OCT signals at the photoreceptor OS tips with the complex conjugate of OCT signals at the IS/OS. The argument of the resulting product yielded temporal phase changes between the two boundaries of the photoreceptor OS. We computed ORG responses for each photoreceptor by integrating phase across a 3 × 3 pixel square centered at the manually selected photoreceptor position and averaging them across up to five repeated imaging sessions. For phase responses exceeding ±*π* radians, the phase was unwrapped along the temporal dimension. The phase change was converted into ΔOPL by multiplying it by *λ*_5_⁄4*π*, where *λ*_5_is the center wavelength of the OCT system, equal to 820nm.

To explore the ORG responses at a higher temporal resolution, we leveraged the scan dimension as a secondary, higher-resolution time axis^13^. The entire volume was divided into five sub-volumes along the scan direction. ORG responses were extracted from individual rods within each sub-volume and averaged to obtain one value for each sub-volume. We assigned the midpoint of the scan-time across individual sub-volumes as the timestamp of the averaged ORG responses. Therefore, a 41.4 Hz volume rate was effectively increased 5-fold in its temporal sampling to reveal features that may otherwise be obfuscated.

## Acknowledgments

We thank Drs. D. Palanker, E. Pugh, J. Jonas, A. Roorda for fruitful discussions. T. Ling acknowledges the support of the National Research Foundation, Singapore for the NRF Fellowship Award (NRF-NRFF14-2022-0005), the Startup Grant from Nanyang Technological University, the seed funding programme under NMRC Centre Grant – Singapore Imaging Eye Network (SIENA) (NMRC/CG/C010A/2017), the National Medical Research Council (MOH-001748-00), and the Ministry of Education, Singapore under its AcRF Tier 1 Grant (RS19/20; RG28/21). R. Sabesan acknowledges support from National Institutes of Health grants U01EY032055, EY029710, unrestricted grant from the Research to Prevent Blindness; George and Martina Kren Professorship in Vision Research, Dawn’s Light Foundation. L. Schmetterer acknowledges support from National Medical Research Council (MOH-000500-00; MOH-000647-00; MOH-000707-00; MOH-001001-00; MOH 001015-00; MOH-001072-06; MOH-001286-00; MOH-001574-02; MOH-001576-00; MOH-001745-00), National Research Foundation Singapore (NRF-CRP24-2020-0001), National Health Innovation Centre Singapore (NHIC-I2D-2402329), SingHealth & Duke-NUS (AM/TP085/2024 [SRDUKAMR2485]; AM/AIR017/2024 [SRDUKAMR24A7]), Duke-NUS (05/FY2022/EX/66-A128; Duke-NUS-KBrFA/2024/0088), Singapore Eye Research Institute & Nanyang Technological University (SERI-NTU Advanced Ocular Engineering (STANCE) Program), and Singapore Eye Research Institute (I010/2024 [2082/44/2024]). V. A. Barathi acknowledges the support for Singapore NMRC Centre Grant (NMRC/CG/M010/2017/Pre-Clinical; NMRC/CG21APR1010-SAMURAI/Pre-Clinical Core Platform/2021_SERI).

## Disclosures

H.L., B.T., L.S. and T.Ling are inventors on a PCT patent application (PCT/SG2024/050050) related to unsupervised signal classification and processing for optoretinography. V.P.P. and R.S. have filed a patent on the line-scan OCT technology used for optoretinography (PCT/US2020/029984).

## Data availability

All data needed to evaluate the conclusions in the paper are present in the paper. Additional data related to this paper may be requested from the authors.

## Author contributions

T.Ling, H.L. and R.S. conceived the study. B.T. and L.S. built the point-scan OCT setup. V.P.P. and R.S. built the line-scan OCT setup. H.L., V.P.P., D.N., C.E.W, T.Liu and P.W.K. conducted the experiments. V.A.B. supported the animal preparation. H.L. and C.E.W. analyzed the data. H.L., C.E.W., R.S. and T.Ling wrote the draft. T.Ling, L.S. and R.S. obtained funding. All work was supervised by T.Ling and R.S. All authors contributed to the final manuscript.

## References

1 Hofmann, K. P. & Lamb, T. D. Rhodopsin, light-sensor of vision. Prog. Retin. Eye Res. 93, 101116 (2023). 10.1016/j.preteyeres.2022.101116

2 Arshavsky, V. Y., Lamb, T. D. & Edward N. Pugh, J. G Proteins and Phototransduction. Annu. Rev. Physiol. 64, 153–187 (2002). 10.1146/annurev.physiol.64.082701.102229

3 Hecht, S., Shlaer, S. & Pirenne, M. H. ENERGY, QUANTA, AND VISION. J. Gen. Physiol. 25, 819–840 (1942). 10.1085/jgp.25.6.819

4 Baylor, D. A., Lamb, T. D. & Yau, K. W. Responses of retinal rods to single photons. The Journal of Physiology 288, 613–634 (1979). 10.1113/jphysiol.1979.sp012716

5 Jackson, G. R., Owsley, C. & Curcio, C. A. Photoreceptor degeneration and dysfunction in aging and age-related maculopathy. Ageing Research Reviews 1, 381–396 (2002). 10.1016/S1568-1637(02)00007-7

6 Curcio, C. A., Medeiros, N. E. & Millican, C. L. Photoreceptor loss in age-related macular degeneration. Invest. Ophthalmol. Vis. Sci. 37, 1236–1249 (1996).

7 Portera-Cailliau, C., Sung, C. H., Nathans, J. & Adler, R. Apoptotic photoreceptor cell death in mouse models of retinitis pigmentosa. Proceedings of the National Academy of Sciences 91, 974–978 (1994). doi:10.1073/pnas.91.3.974

8 Jackson, G. R. & Owsley, C. Scotopic sensitivity during adulthood. Vision Res. 40, 2467–2473 (2000). 10.1016/S0042-6989(00)00108-5

9 Owsley, C. et al. Delayed Rod-Mediated Dark Adaptation Is a Functional Biomarker for Incident Early Age-Related Macular Degeneration. Ophthalmology 123, 344–351 (2016). 10.1016/j.ophtha.2015.09.041

10 Saszik, S. M., Robson, J. G. & Frishman, L. J. The Scotopic Threshold Response of the Dark-Adapted Electroretinogram of the Mouse. The Journal of Physiology 543, 899–916 (2002). 10.1113/jphysiol.2002.019703

11 Lamb, T. D. & Pugh, E. N. Dark adaptation and the retinoid cycle of vision. Prog. Retin. Eye Res. 23, 307–380 (2004). 10.1016/j.preteyeres.2004.03.001

12 Lamb, T. D. & Pugh, E. N., Jr. Phototransduction, Dark Adaptation, and Rhodopsin Regeneration The Proctor Lecture. Invest. Ophthalmol. Vis. Sci. 47, 5138–5152 (2006). 10.1167/iovs.06-0849

13 Pandiyan, V. P. et al. The optoretinogram reveals the primary steps of phototransduction in the living human eye. Science Advances 6, eabc1124 (2020).

14 Tomczewski, S. et al. Photopic flicker optoretinography captures the light-driven length modulation of photoreceptors during phototransduction. Proceedings of the National Academy of Sciences 122, e2421722122 (2025). doi:10.1073/pnas.2421722122

15 Kim, T.-H., Ding, J. & Yao, X. Intrinsic signal optoretinography of dark adaptation kinetics. Sci. Rep. 12, 2475 (2022). 10.1038/s41598-022-06562-4

16 Pandiyan, V. P., Nguyen, P. T., Pugh, E. N. & Sabesan, R. Human cone elongation responses can be explained by photoactivated cone opsin and membrane swelling and osmotic response to phosphate produced by RGS9-catalyzed GTPase. Proceedings of the National Academy of Sciences 119, e2202485119 (2022). doi:10.1073/pnas.2202485119

17 Hillmann, D. et al. In vivo optical imaging of physiological responses to photostimulation in human photoreceptors. Proc. Natl. Acad. Sci. U. S. A. 113, 13138–13143 (2016). 10.1073/pnas.1606428113

18 Zhang, F., Kurokawa, K., Lassoued, A., Crowell, J. A. & Miller, D. T. Cone photoreceptor classification in the living human eye from photostimulation-induced phase dynamics. Proc. Natl. Acad. Sci. U. S. A. 116, 7951–7956 (2019). 10.1073/pnas.1816360116

19 Azimipour, M. et al. Optoretinogram: optical measurement of human cone and rod photoreceptor responses to light. Opt Lett 45, 4658–4661 (2020). 10.1364/OL.398868

20 Lassoued, A. et al. Cone photoreceptor dysfunction in retinitis pigmentosa revealed by optoretinography. Proceedings of the National Academy of Sciences 118, e2107444118 (2021). doi:10.1073/pnas.2107444118

21 Tomczewski, S. et al. Light-adapted flicker optoretinograms captured with a spatio-temporal optical coherence-tomography (STOC-T) system. Biomedical Optics Express 13, 2186–2201 (2022). 10.1364/BOE.444567

22 Tan, B. et al. Light-evoked deformations in rod photoreceptors, pigment epithelium and subretinal space revealed by prolonged and multilayered optoretinography. Nature Communications 15, 5156 (2024). 10.1038/s41467-024-49014-5

23 Zhang, P. et al. In vivo optophysiology reveals that G-protein activation triggers osmotic swelling and increased light scattering of rod photoreceptors. Proc. Natl. Acad. Sci. U. S. A. 114, E2937–E2946 (2017). 10.1073/pnas.1620572114

24 Pandiyan, V. P. et al. Characterizing cone spectral classification by optoretinography. Biomedical Optics Express 13, 6574–6594 (2022). 10.1364/BOE.473608

25 Boyle, K. C. et al. Mechanisms of Light-Induced Deformations in Photoreceptors. Biophys. J. 119, 1481–1488 (2020). 10.1016/j.bpj.2020.09.005

26 Mustafi, D., Engel, A. H. & Palczewski, K. Structure of cone photoreceptors. Prog. Retin. Eye Res. 28, 289–302 (2009). 10.1016/j.preteyeres.2009.05.003

27 Volland, S. et al. Three-dimensional organization of nascent rod outer segment disk membranes. Proceedings of the National Academy of Sciences 112, 14870–14875 (2015). doi:10.1073/pnas.1516309112

28 Makino, C. L., Taylor, W. R. & Baylor, D. A. Rapid charge movements and photosensitivity of visual pigments in salamander rods and cones. The Journal of Physiology 442, 761–780 (1991). 10.1113/jphysiol.1991.sp018818

29 Kessler, C., Tillman, M., Burns, M. E. & Pugh Jr, E. N. Rhodopsin in the rod surface membrane regenerates more rapidly than bulk rhodopsin in the disc membranes in vivo. The Journal of Physiology 592, 2785–2797 (2014). 10.1113/jphysiol.2014.272518

30 Berger, A. et al. Spectral-Domain Optical Coherence Tomography of the Rodent Eye: Highlighting Layers of the Outer Retina Using Signal Averaging and Comparison with Histology. PLoS One 9, e96494 (2014). 10.1371/journal.pone.0096494

31 Guziewicz, K. E. et al. BEST1 gene therapy corrects a diffuse retina-wide microdetachment modulated by light exposure. Proceedings of the National Academy of Sciences 115, E2839 (2018). 10.1073/pnas.1720662115

32 Lindell, M. et al. Volumetric Reconstruction of a Human Retinal Pigment Epithelial Cell Reveals Specialized Membranes and Polarized Distribution of Organelles. Invest. Ophthalmol. Vis. Sci. 64, 35–35 (2023). 10.1167/iovs.64.15.35

33 Preston, G. M., Carroll, T. P., Guggino, W. B. & Agre, P. Appearance of Water Channels in Xenopus Oocytes Expressing Red Cell CHIP28 Protein. Science 256, 385–387 (1992). doi:10.1126/science.256.5055.385

34 Brown, K. T. & Murakami, M. A New Receptor Potential of the Monkey Retina with no Detectable Latency. Nature 201, 626–628 (1964). 10.1038/201626a0

35 Cone, R. A. Early Receptor Potential of the Vertebrate Retina. Nature 204, 736–739 (1964). 10.1038/204736a0

36 Hodgkin, A. L. & Obryan, P. M. Internal recording of the early receptor potential in turtle cones. The Journal of Physiology 267, 737–766 (1977). 10.1113/jphysiol.1977.sp011836

37 Murakami, M. & Pak, W. L. Intracellularly recorded early receptor potential of the vertebrate photoreceptors. Vision Res. 10, 965–975 (1970). 10.1016/0042-6989(70)90074-X

38 Ling, T. et al. High-speed interferometric imaging reveals dynamics of neuronal deformation during the action potential. Proceedings of the National Academy of Sciences 117, 10278–10285 (2020). doi:10.1073/pnas.1920039117

39 Yang, Y. et al. Imaging Action Potential in Single Mammalian Neurons by Tracking the Accompanying Sub-Nanometer Mechanical Motion. ACS Nano 12, 4186–4193 (2018). 10.1021/acsnano.8b00867

40 You, H., Li, H. & Ling, T. Electromechanical deformation of biological neurons: an intrinsic marker for label-free functional neuroimaging. Journal of Physics D: Applied Physics (2024).

41 Pak, W. L. & Cone, R. A. Isolation and Identification of the Initial Peak of the Early Receptor Potential. Nature 204, 836–838 (1964). 10.1038/204836a0

42 Marsh, D. Renormalization of the tension and area expansion modulus in fluid membranes. Biophys. J. 73, 865–869 (1997). 10.1016/S0006-3495(97)78119-0

43 Pandiyan, V. P., Jiang, X., Kuchenbecker, J. A. & Sabesan, R. Reflective mirror-based line-scan adaptive optics OCT for imaging retinal structure and function. Biomedical Optics Express 12, 5865–5880 (2021). 10.1364/BOE.436337

44 Liebman, P. A., Jagger, W. S., Kaplan, M. W. & Bargoot, F. G. Membrane structure changes in rod outer segments associated with rhodopsin bleaching. Nature 251, 31–36 (1974). 10.1038/251031a0

45 Kaplan, M. W. Modeling the rod outer segment birefringence change correlated with metarhodopsin II formation. Biophys. J. 38, 237–241 (1982). 10.1016/S0006-3495(82)84554-2

46 Kühn, H., Bennett, N., Michel-Villaz, M. & Chabre, M. Interactions between photoexcited rhodopsin and GTP-binding protein: kinetic and stoichiometric analyses from light-scattering changes. Proceedings of the National Academy of Sciences 78, 6873–6877 (1981). 10.1073/pnas.78.11.6873

47 Hofmann, K. P., Schleicher, A., Emeis, D. & Reichert, J. Light-induced axial and radial shrinkage effects and changes of the refractive index in isolated bovine rod outer segments and disc vesicles. Biophys. Struct. Mech. 8, 67–93 (1981). 10.1007/BF01047107

48 Michel-Villaz, M., Brisson, A., Chapron, Y. & Saibil, H. Physical analysis of light-scattering changes in bovine photoreceptor membrane suspensions. Biophys. J. 46, 655–662 (1984). 10.1016/S0006-3495(84)84064-3

49 Jagger, W. S. & Liebman, P. A. Anomalous dispersion of rhodopsin in rod outer segments of the frog. J. Opt. Soc. Am. 66, 56–59 (1976). 10.1364/JOSA.66.000056

50 Chabre, M. & Cavaggioni, A. Light Induced Changes of Ionic Flux in the Retinal Rod. Nature New Biology 244, 118–120 (1973). 10.1038/newbio244118a0

51 Yagi, N. Structural changes in rod outer segments of frog and mouse after illumination. Exp. Eye Res. 116, 395–401 (2013). 10.1016/j.exer.2013.09.016

52 Fu, Y. & Yau, K.-W. Phototransduction in mouse rods and cones. Pflügers Archiv - European Journal of Physiology 454, 805–819 (2007). 10.1007/s00424-006-0194-y

53 Hendrickson, A. & Drucker, D. The development of parafoveal and mid-peripheral human retina. Behav. Brain Res. 49, 21–31 (1992). 10.1016/S0166-4328(05)80191-3

54 Goldstein, E. B. & Berson, E. L. Cone Dominance of the Human Early Receptor Potential. Nature 222, 1272–1273 (1969). 10.1038/2221272a0

55 Cone, R. A. et al. The early receptor potential. principles of receptor physiology, 345–365 (1971).

56 Campbell, M. C. & Hughes, A. An analytic, gradient index schematic lens and eye for the rat which predicts aberrations for finite pupils. Vision Res. 21, 1129–1148 (1981). 10.1016/0042-6989(81)90016-X

57 Perlman, I. Kinetics of bleaching and regeneration of rhodopsin in abnormal (RCS) and normal albino rats in vivo. The Journal of Physiology 278, 141–159 (1978). 10.1113/jphysiol.1978.sp012297

58 Zhang, P., Goswami, M., Zawadzki, R. J. & Pugh, E. N., Jr. The Photosensitivity of Rhodopsin Bleaching and Light-Induced Increases of Fundus Reflectance in Mice Measured In Vivo With Scanning Laser Ophthalmoscopy. Invest. Ophthalmol. Vis. Sci. 57, 3650–3664 (2016). 10.1167/iovs.16-19393

59 Li, H. et al. Phase-restoring subpixel image registration: enhancing motion detection performance in Fourier-domain optical coherence tomography. Journal of Physics D: Applied Physics 58, 145102 (2025). 10.1088/1361-6463/adb3b4

60 Guizar-Sicairos, M., Thurman, S. T. & Fienup, J. R. Efficient subpixel image registration algorithms. Optics letters 33, 156–158 (2008).

61 Spahr, H. et al. Phase-sensitive interferometry of decorrelated speckle patterns. Sci. Rep. 9, 11748 (2019). 10.1038/s41598-019-47979-8

62 Stevenson, S. & Roorda, A. Correcting for miniature eye movements in high-resolution scanning laser ophthalmoscopy. Vol. 5688 PWB (SPIE, 2005).

