## Supplementary Information for "Optoretinography reveals rapid rod photoreceptor movement upon photoisomerization"

### Supplementary Section 1: Separating the light-evoked dynamics of rod OS tips and the RPE-BrM complex using unsupervised learning

We applied principal component analysis across all phase traces in Fig. 1B and projected them onto the first principal component, which captured 98.5% of the total variance (see Supplementary Fig. 1A). After this dimensionality reduction, the distribution of the principal component score (orange bins) exhibited a bimodal distribution and was fitted into a Gaussian mixture model (GMM) with two components. The dashed magenta line in Supplementary Fig. 1A denotes the mixture distribution, and the gray curves represent two individual components. Two signal patterns were clustered into Type-I and Type-II based on the posterior probability, as indicated by the Bayes boundary drawn in Supplementary Fig. 1A.

To consistently separate the two light-evoked dynamic patterns across measurements at different stimulus strengths, we analyzed the distribution of two signal patterns in space. For each data point in Supplementary Fig. 1A, we retrieved its spatial location, calculated the distance to the bottom of the BrM, and further obtained a normalized distance that was divided by the mean thickness of the composite layer. Supplementary Fig. 2A illustrates the distribution of the normalized distances for the two types of signals. Based on this depth distribution profile, Type-I corresponds to rod OS tips, while Type-II can be attributed to the RPE-BrM complex[1]. We explored optimal decision boundaries for isolating these retinal structures based on the $F_\beta$ score, which is defined as,

$$F_\beta = (1+\beta^2)\cdot\frac{\text{precision}\cdot\text{recall}}{\beta^2\cdot\text{precision}+\text{recall}}, \tag{S1}$$

$$\text{precision} = \frac{TP}{TP+FP}, \tag{S2}$$

$$\text{recall} = \frac{TP}{TP+FN}, \tag{S3}$$

where $TP$, $FN$, and $FP$ represent the number of true positives, false negatives, and false positives, respectively.

In this study, we set $\beta$ to 0.5 to place more emphasis on precision over recall. Supplementary Fig. 2B shows the $F_{0.5}$ scores calculated for both signal types at varying thresholds. For each signal type, the optimal threshold was determined based on the depth that achieves the maximum $F_{0.5}$ score (see red and blue stars in Supplementary Fig. 2B). The associated normalized confusion matrices were shown in Supplementary Figs. 2C-D. Accordingly, the composite layer can be divided into rod OS tips, a transition region, and the RPE-BrM complex.

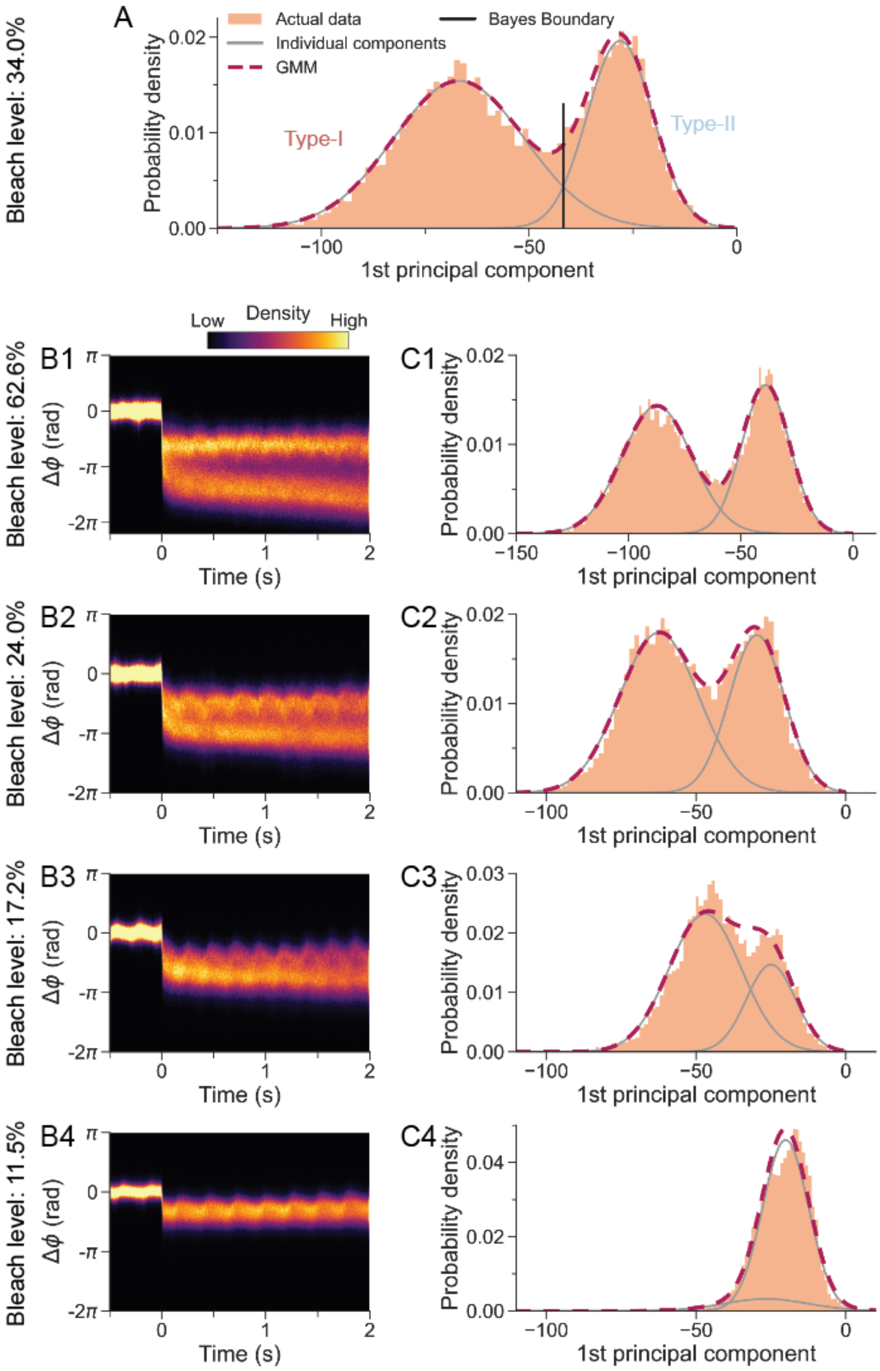


**Supplementary Figure 1.** Distributions of phase traces evoked by bleaching stimuli at varying bleach levels. (A) Each phase trace in Fig. 1B was projected onto the 1st principal component. The distribution of the principal component score (orange bins) was fitted into a Gaussian mixture model (GMM, dashed magenta line). The gray curves represent two individual components, and the dashed magenta line denotes the mixture distribution. Two types of signals were separated according to the Bayes boundary (solid black line). (B) Distribution density maps of phase traces extracted from the composite layer using the inner segment/outer segment junction as the reference. (C) Principal component analysis was independently conducted on each dataset, and phase traces were projected onto the corresponding 1st principal component.

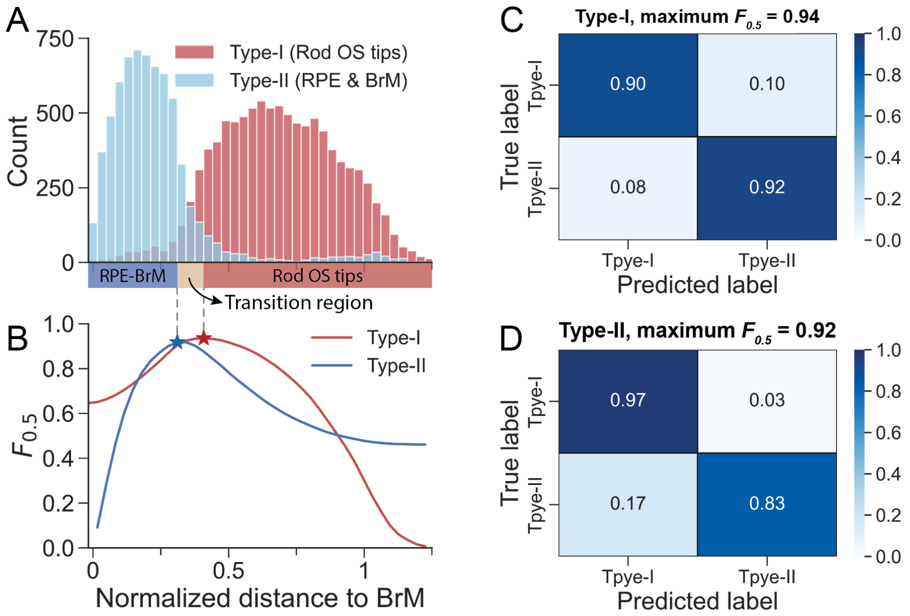


**Supplementary Figure 2.** Clustering of two signal patterns based on the axial distance to Bruch's membrane (BrM). (A) The depth distribution of signals shown in Supplementary Fig. 1A. For each pixel, the distance to BrM was calculated and normalized by dividing the average thickness of the composite layer. (B) $F_{0.5}$ scores for the Type-I signal (red curve) and Type-II signal (blue curve) at varying depth thresholds. Star markers indicate the thresholds with the highest $F_{0.5}$ scores. Normalized confusion matrices at the thresholds indicated by the (C) red star and (D) blue star in panel B.

### Supplementary Section 2: Testing whether the rapid OS shrinkage can be explained based on the osmolarity-driven water efflux model

If the rapid shrinkage of the rod OS is driven by a sudden decrease in osmotic pressure, the water permeability coefficient of the rod OS plasma membrane can be estimated based on a model developed by Zhang *et al.*[2], with parameters listed in Supplementary Table 1. Assuming that the shrinkage is purely an osmotic effect, Van't Hoff's law suggests that the saturated decrease in cytoplasmic volume, denoted as $\Delta V_{\mathrm{cyto}}$, is proportional to the initial decrease in the osmotic pressure,

$$\Delta V_{\mathrm{cyto}}/V_{\mathrm{cyto,\,rest}} = \Delta \Pi/\Pi_{\mathrm{rest}}, \tag{S4}$$

where $V_{\mathrm{cyto,\,rest}}$ and $\Pi_{\mathrm{rest}}$ represent the cytoplasmic volume and osmotic pressure of the rod OS in the rest (dark-adapted) state. $\Delta \Pi$ is the change in the osmatic pressure triggered by visual stimuli. Since moderate osmotic changes do not affect the width of the rod OS[2] and the cytoplasmic space occupies about 44% of the space within the rod OS[3], the fractional volume decrease $\Delta V_{\mathrm{cyto}}/V_{\mathrm{cyto,\,rest}}$ can be calculated as $\Delta L_{\mathrm{ROS}}/(0.44 \times L_{\mathrm{ROS}})$. In response to a flash at 62.6% bleach, the change in cytoplasmic pressure can be calculated by $\Delta \Pi/RT = [\Delta L_{\mathrm{ROS}}/(0.44 \times L_{\mathrm{ROS}})] \cdot (\Pi_{\mathrm{rest}}/RT) = -4.39\ \mathrm{mOsM}$.

The water flow rate depends on the osmotic pressure difference across the membrane and the hydraulic conductivity $L_p$ of the plasma membrane. The initial water flow rate following a sudden decrease in the osmotic pressure, denoted as $J_w$, can be calculated by,

$$J_w = L_p S_{\mathrm{ROS}} \Delta \Pi, \tag{S5}$$

In our measurements, we found that the rapid contraction saturated 10 ms after the flash onset. Since the water flow rate driven by a sudden osmolarity change would gradually decrease as it approaches a new osmotic equilibrium, a conservative estimation of the initial flow rate is $\Delta L_{\mathrm{ROS}}/10\ \mathrm{ms} = -14.26\ \mu\mathrm{m/s}$, or $-32.37\ \mu\mathrm{m}^3/\mathrm{s}$ with a cross-sectional area of $2.27\ \mu\mathrm{m}^2$. According to Eq. (S5), the hydraulic conductivity $L_p$ can be calculated as $216.78\ \mu\mathrm{m}^3/(\mathrm{s} \cdot \mathrm{dyn})$, and the corresponding water permeability coefficient is $0.30\ \mathrm{cm/s}$. This value is much larger than $2.6 \times 10^{-3}\ \mathrm{cm/s}$ measured in previous in-vitro studies[4]. This discrepancy suggests that the observed rapid shrinkage of the rod OS is too fast to be attributed to osmosis.

**Supplementary Table 1.** Parameters used for testing the osmolarity-driven water efflux model

| Parameter | Value | Unit |
|---|---|---|
| Normal rodent plasma osmolarity[5], $\Pi_{\mathrm{rest}}/RT$ | 325 | mOsM |
| Length of the rod OS[6], $L_{\mathrm{ROS}}$ | 24.0 | µm |
| Maximum contraction*, $\Delta L_{\mathrm{ROS}}$ | -142.6 | nm |
| Surface area of the rod outer segment, $S_{\mathrm{ROS}}$ | 132 | $\mu\mathrm{m}^2$ |
| Cross-sectional area of the rod OS[6], $A_{\mathrm{ROS}}$ | 2.27 | $\mu\mathrm{m}^2$ |
| Initial water influx rate*, $J_w$ | -32.37 | $\mu\mathrm{m}^3/\mathrm{s}$ |

* Calculated with a refractive index of 1.41.

### Supplementary Section 3: Contraction of the rod OS based on the voltage-dependent membrane tension model

The disk membranes in cones are confluent with the plasma membrane, while most rod disk membranes, except for a few nascent basal disks, are entirely enclosed by the plasma membrane[7,8]. Despite the distinct membranous organization of cone and rod OS[9], the following discussion provides justifications for applying the electromechanical coupling model (Eq. 1), which was established to explain the rapid contraction in the cone OS, to characterize the contraction in the rod OS.

**1) The electromechanical coupling model proposed for cone early receptor potential (ERP) primarily concerns the shape changes of individual disks, which would apply to the disks in rods as well.** Although rods and cones are drastically different in their membrane organization, they can share similar changes in voltage-dependent membrane tension during the ERP in individual disks. Such a driving force of voltage-dependent membrane tension can be used to estimate the steady-state equilibrium of individual disk area expansion/contraction without considering other properties of the rod OS, such as the viscosity of the cytoplasm, that may affect the time-dependent dynamics.

**2) The slow recovery from rapid contraction enables modeling the rod ERP quasi-statically.** In response to strong bleaching stimuli, measured ΔOPL in rodent rod OS exhibited a persistent contraction. The slow recovery rate makes the quasi-static model suitable for calculating the amplitude of the rapid contraction response in the rod OS.

Given the above considerations, we adopted the quasi-static electromechanical coupling model based on the voltage-dependent membrane tension during the ERP[10] (see Eq. 1) to model the rapid rod OS contraction, particularly its amplitude, with parameters listed in Table 1 and Supplementary Table S2.

To link the area expansion in Eq. 1 with the measured ΔOPL, we assume a constant volume of individual disks during the millisecond-scale dynamics. The axial shrinkage of each disk caused by its area expansion can thus be estimated by[10],

$$\Delta z = -\frac{\Delta A}{A} z, \tag{S6}$$

For the entire photoreceptor OS, we can calculate ΔOPL as,

$$\Delta \mathrm{OPL} = \Delta z \times N_{\mathrm{disks}} \times n_{\mathrm{index}}, \tag{S7}$$

where $N_{\mathrm{disks}}$ is the number of disks along each OS, $n_{\mathrm{index}}$ is the refractive index.

**Supplementary Table 2.** Parameters used for calculating the rod OS contraction based on the voltage-dependent membrane tension model

| Parameter | Value | Unit |
| --- | --- | --- |
| Length of the rod OS[6], $L_{\mathrm{ROS}}$ | 24.0 | μm |
| Cross-sectional area of disk face[6], $A$ | 2.1 | $\mu m^2$ |
| Disc-disc distance[3], $d$ | 32 | nm |
| Number of disks per rod, $N_{\mathrm{disks}} = L_{\mathrm{ROS}}/d$ | 750 | / |
| Height of a single disk[3], $z$ | 18 | nm |
| Rhodopsin density on the disc membrane[11], $\sigma_{\mathrm{rho}}$ | 23000 | $\mathrm{molecules}/\mu\mathrm{m}^2$ |
| Charge displacement per photoisomerization[*], $q$ | 0.14 | electron |
| Specific membrane capacitance[12,13], $c_m$ | 1 | $\mu\mathrm{F}/\mathrm{cm}^2$ |
| Area expansion modulus[10], $K_A$ | 0.2 | $\mathrm{N}/\mathrm{m}$ |
| Tension/voltage[10,14] | -0.1 | $\mathrm{mN}/(\mathrm{m}{\cdot}\mathrm{V})$ |

* Charge displacement per photoisomerization was estimated as follows:

Previous studies have suggested that a full bleach results in the displacement of $2 \times 10^5$ electronic charges across the rod plasma membrane[13], or equivalently $6.2 \times 10^4$ electronic charges shift at a 30% bleach level[15]. Considering that each rod contains $7 \times 10^7$ rhodopsin molecules[11], and 2% of rhodopsin molecules are located in the plasma membrane and nascent basal disks[16], the charge displacement per photoisomerization can be calculated as $2 \times 10^5/(7 \times 10^7 \times 2\%) = 0.14$ electrons.

† Voltage change at the disk membranes can be calculated by $\Delta V_m = \text{bleach level} \times \sigma_{rho} \times q\ /c_m$.

### Supplementary Section 4: Calculation of bleach levels

#### 4.1 Rhodopsin bleaching in rodents

Given the negligible regeneration of rhodopsin during visual stimulation, rhodopsin bleaching can be calculated by[17,18],

$$p(p_0, Q) = p_0 \exp(-Q/Q_e), \tag{S8}$$

$$\text{Bleach level} = p_0 - p(p_0, Q) = p_0[1 - \exp(-Q/Q_e)], \tag{S9}$$

where $p_0$ and $p(p_0, Q)$ represent the fraction of rhodopsin present before and after visual stimulation, respectively. We set $p_0$ to 1 when calculating the rhodopsin bleaching in rats, as the animals were dark-adapted overnight before the ORG experiments[17]. $Q$ is the effective energy density (photons/μm$^2$) of the visual stimulus, which can be obtained by[18],

$$Q = \frac{t_f}{A} \cdot \int_0^{+\infty} \frac{P(\lambda)S(\lambda)}{E_v(\lambda)} d\lambda, \tag{S10}$$

where $t_f$ is the duration of the flash, $A$ is the illuminated area on the retina, $P(\lambda)$ is the power spectral density of the visual stimulus (W/nm), $S(\lambda)$ is the normalized absorption spectrum of the visual pigment calculated based on the Lamb template[19]. $E_v(\lambda)$ is the energy of a single photon at wavelength $\lambda$ and can be calculated by $hc/\lambda$, where $h$ and $c$ represent the Planck constant and the speed of light in vacuum, respectively. $Q_e$ is the energy density that reduces rhodopsin to $1/\mathrm{e}$ of its dark-adapted level, which was measured to be 7.94×10$^7$ photons/μm$^2$ in rats[17]. Note that both $Q$ and $Q_e$ are calculated by dividing the energy measured at the cornea by the illumination area on the retina[18]. The bleach levels of flashes used in rodent ORG experiments are summarized in Supplementary Table 3.

**Supplementary Table 3.** Stimulus parameters in rodent ORG experiments.

| **Energy density $Q$ (photons/μm$^2$)** | **Bleach level (%)** |
|---|---|
| $1.88 \times 10^6$ | 2.3 |
| $2.83 \times 10^6$ | 3.5 |
| $4.27 \times 10^6$ | 5.2 |
| $6.72 \times 10^6$ | 8.1 |
| $9.75 \times 10^6$ | 11.5 |
| $1.50 \times 10^7$ | 17.2 |
| $2.17 \times 10^7$ | 24.0 |
| $3.30 \times 10^7$ | 34.0 |
| $5.04 \times 10^7$ | 47.0 |
| $7.81 \times 10^7$ | 62.6 |

### 4.2 Calculation of bleach levels in human subjects

The bleach level in human rods can be calculated by[20,21],

$$\text{Bleach level} = p_0\left[1 - \exp\left(-Q_{\text{retina}}/Q_{e,\text{retina}}\right)\right], \quad \text{(S11)}$$

where $Q_{\text{retina}}$ is the energy density of the flash stimulus at the retinal surface, which can be calculated by,

$$Q_{\text{retina}} = \frac{t_f}{A} \cdot \int_0^{+\infty} \frac{\tau(\lambda)P(\lambda)S(\lambda)}{E_v(\lambda)} d\lambda, \quad \text{(S12)}$$

Note that compared to Eq. (S10), the transmissivity of the ocular media[22], denoted by $\tau(\lambda)$, is incorporated into the calculation in Eq. (S12).

For rhodopsin bleaching, $Q_{e,\text{retina}}$, which represents the energy density required to reduce rhodopsin to $1/\text{e}$ of its dark-adapted level, can be calculated by,

$$Q_{e,\text{retina}} = 1/f_{\text{Dens}} f_{wg} \alpha_{\lambda_{\text{max}}} \gamma, \quad \text{(S13)}$$

where $f_{\text{Dens}}$ is a factor accounting for the "self-screening" effect, i.e., the decrease in the power along the length of the dark-adapted rod OS[20,21]. $f_{\text{Dens}}$ has been calculated to be $1/1.44$, given an axial density of 0.45 in the human rod OS[20]. $f_{wg}\alpha_{\lambda_{\text{max}}}\gamma$ represents rhodopsin photosensitivity and has been measured to be $3.1 \times 10^{-8}$ $\mu\text{m}^2/\text{photon}$ using a scanning laser ophthalmoscope[20] in humans.

The fraction of rhodopsin present after a 15-min dark adaptation, can be calculated according to the "MLP rate-limited kinetic model"[20],

$$p_0 = 1 - K_m W\left\{\frac{B_0}{K_m}\exp\left(\frac{B_0}{K_m}\right)\exp\left(-\frac{1+K_m}{K_m}rt\right)\right\}, \quad \text{(S14)}$$

where $W(x)$ is the "Lambert W" function[23]. $B_0$ represents the fraction of rhodopsin bleached before dark adaptation. The parameters $K_m$ and $r$ were 0.25 and 0.8 $\text{min}^{-1}$, respectively, based on the time course of rhodopsin regeneration measured in normal human subjects[20]. Using $B_0 = 1$ and $t = 15$ min, we obtained $p_0 = 0.91$, indicating that a 15-min dark adaptation following full bleaching regenerates 91% of rhodopsin. The energy densities and the corresponding bleach levels of flashes used in human ORG experiments are summarized in Supplementary Table 4.

**Supplementary Table 4.** Stimulus parameters in human rod ORG experiments.

| Energy density $Q_{\text{retina}}$ (photons/μm$^2$) | Bleach level (%) |
|---|---|
| $2.83 \times 10^5$ | 0.6 |
| $1.70 \times 10^6$ | 3.3 |
| $7.79 \times 10^6$ | 14.0 |
| $1.65 \times 10^7$ | 27.1 |
| $2.65 \times 10^7$ | 39.4 |

In 1955, Hagins reported that a brief flash (< 1 ms), no matter how strong it is, cannot bleach more than 50% of rhodopsin in the living rabbit eye[24]. This observation was later explained by the ability of early photoproducts, up to Meta I, to absorb a second photon and revert back (photoreversal) to rhodopsin or its isoform (isorhodopsin)[25,26]. Given the millisecond-scale thermal decay time constant of Meta I[27], this effect is expected to be marginal in our current experiments using 5-ms flashes. Nonetheless, a future study that calculates the bleach level considering the entire rhodopsin cycle would be valuable.

### Supplementary Figures

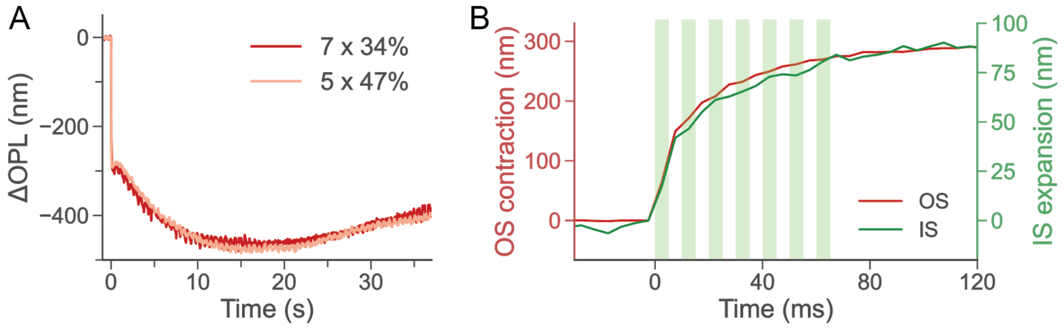


**Supplementary Figure 3.** ORG responses in the rod IS and OS evoked by serial flashes with a flash duration of 5 ms and an inter-flash interval of 5 ms. (A) Rod OS responses evoked by two sets of serial-flash stimuli: one with seven flashes, each at a 34% bleach level (red curve), and the other with five flashes, each at a 47% bleach level (pink curve). Both configurations were expected to bleach approximately 95% of the total rhodopsin (94.5% vs 95.8%) by the end of the stimulation process. (B) The rapid OS contraction and the corresponding IS expansion in response to seven flashes, each at a 34% bleach level. The green blocks show the timing of the multiple flash stimuli. The rapid OS responses evoked by each flash in sequence gradually decayed, which followed the expected trend for a smaller number of rhodopsins bleached after each flash. The IS expansion and OS contraction exhibit similar trends.

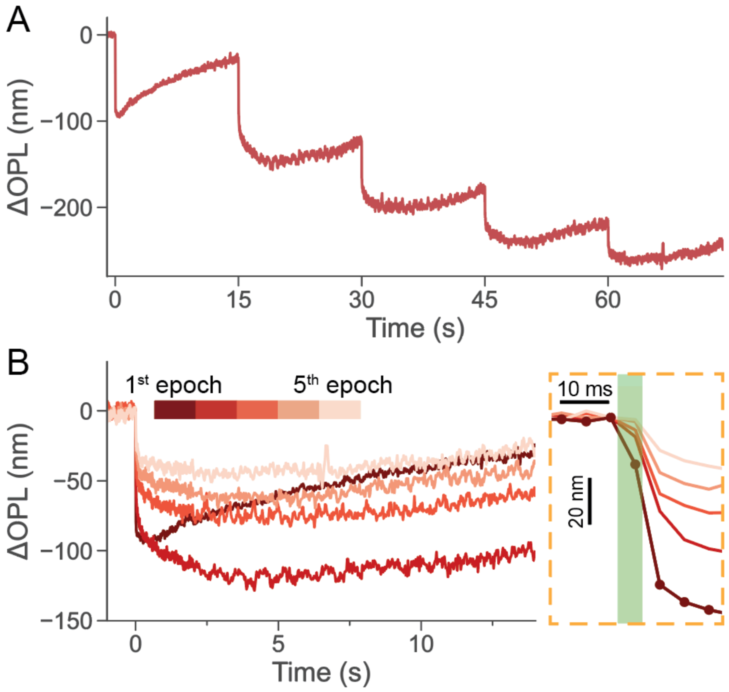


**Supplementary Figure 4.** ORG responses in the rod OS elicited by multiple flashes with an inter-flash interval of 15 s, where each flash bleached 8.1% of the remaining rhodopsins. (A) Prolonged ORG recording of the light-evoked rod OS response during the multi-flash stimulation protocol. (B) The rod OS responses extracted from each epoch were aligned to the flash onsets, with the enlarged view shown in the dashed rectangle.

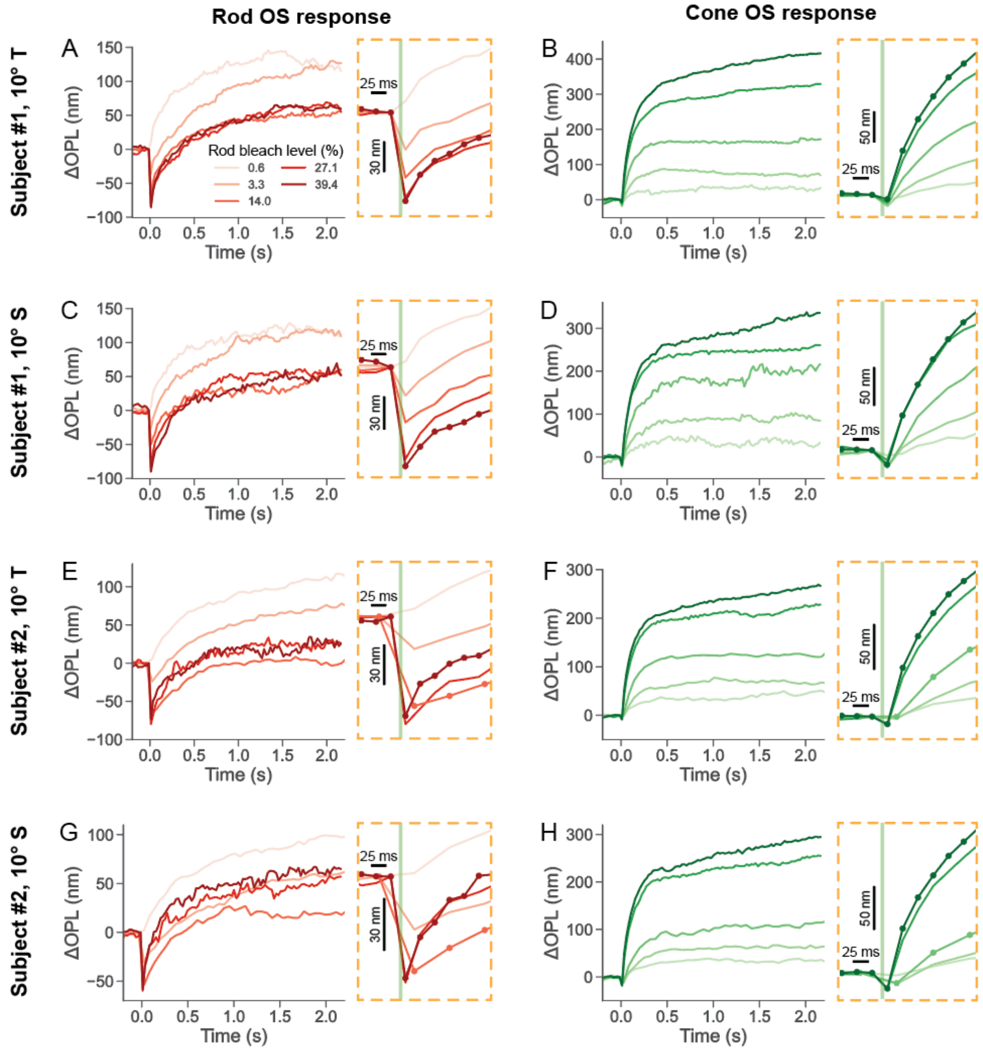


**Supplementary Figure 5.** ORG signals in human rod OS and cone OS. Stimulus strengths ranged from $2.8 \times 10^5 - 2.7 \times 10^7$ photons/$\mu m^2$, corresponding to 0.6% (light color) to 39.4% (deep color) rhodopsin bleach (panel A). Protocol 1, with a volumetric scan rate of 41.4 Hz, was used for the first subject at all bleach levels and for the second subject at 25.0% and 33.9% bleach levels. Protocol 2, with a volumetric scan rate of 17.3 Hz, was used for the second subject at 0.7%, 3.2%, and 12.8% bleach levels.

### Supplementary References


1 Tan, B. *et al.* Light-evoked deformations in rod photoreceptors, pigment epithelium and subretinal space revealed by prolonged and multilayered optoretinography. *bioRxiv*, 2023.2003.2003.530926 (2023). https://doi.org:10.1101/2023.03.03.530926

2 Zhang, P. *et al.* In vivo optophysiology reveals that G-protein activation triggers osmotic swelling and increased light scattering of rod photoreceptors. *Proc. Natl. Acad. Sci. U. S. A.* **114**, E2937-E2946 (2017). https://doi.org:10.1073/pnas.1620572114

3 Pöge, M. *et al.* Determinants shaping the nanoscale architecture of the mouse rod outer segment. *eLife* **10**, e72817 (2021). https://doi.org:10.7554/eLife.72817

4 Preston, G. M., Carroll, T. P., Guggino, W. B. & Agre, P. Appearance of Water Channels in Xenopus Oocytes Expressing Red Cell CHIP28 Protein. *Science* **256**, 385-387 (1992). https://doi.org:doi:10.1126/science.256.5055.385

5 Russell, E. S. & Bernstein, S. E. Blood and blood formation. *Biology of the laboratory mouse* **2**, 351-372 (1966).

6 Hagins, W. A., Penn, R. D. & Yoshikami, S. Dark Current and Photocurrent in Retinal Rods. *Biophys. J.* **10**, 380-412 (1970). https://doi.org:10.1016/S0006-3495(70)86308-1

7 Volland, S. *et al.* Three-dimensional organization of nascent rod outer segment disk membranes. *Proceedings of the National Academy of Sciences* **112**, 14870-14875 (2015). https://doi.org:doi:10.1073/pnas.1516309112

8 Burgoyne, T. *et al.* Rod disc renewal occurs by evagination of the ciliary plasma membrane that makes cadherin-based contacts with the inner segment. *Proceedings of the National Academy of Sciences* **112**, 15922-15927 (2015). https://doi.org:doi:10.1073/pnas.1509285113

9 Mustafi, D., Engel, A. H. & Palczewski, K. Structure of cone photoreceptors. *Prog. Retin. Eye Res.* **28**, 289-302 (2009). https://doi.org:https://doi.org/10.1016/j.preteyeres.2009.05.003

10 Boyle, K. C. *et al.* Mechanisms of Light-Induced Deformations in Photoreceptors. *Biophys. J.* **119**, 1481-1488 (2020). https://doi.org:10.1016/j.bpj.2020.09.005

11 Lyubarsky, A. L., Daniele, L. L. & Pugh, E. N. From candelas to photoisomerizations in the mouse eye by rhodopsin bleaching in situ and the light-rearing dependence of the major components of the mouse ERG. *Vision Res.* **44**, 3235-3251 (2004). https://doi.org:https://doi.org/10.1016/j.visres.2004.09.019

12 Makino, C. L., Taylor, W. R. & Baylor, D. A. Rapid charge movements and photosensitivity of visual pigments in salamander rods and cones. *The Journal of Physiology* **442**, 761-780 (1991). https://doi.org:https://doi.org/10.1113/jphysiol.1991.sp018818

13 Rüppel, H. & Hagins, W. A. in *Biochemistry and Physiology of Visual Pigments.* (ed Helmut Langer) 257-261 (Springer Berlin Heidelberg).

14 Ling, T. *et al.* Full-field interferometric imaging of propagating action potentials. *Light: Science & Applications* **7**, 107 (2018). https://doi.org:10.1038/s41377-018-0107-9

15 Hochstrate, P., Lindau, M. & Rüppel, H. On the Origin and the Signal-Shaping Mechanism of the Fast Photosignal in the Vertebrate Retina. *Biophys. J.* **38**, 53-61 (1982). https://doi.org:10.1016/S0006-3495(82)84530-X

16 Kessler, C., Tillman, M., Burns, M. E. & Pugh Jr, E. N. Rhodopsin in the rod surface membrane regenerates more rapidly than bulk rhodopsin in the disc membranes in vivo. *The Journal of Physiology* **592**, 2785-2797 (2014). https://doi.org:https://doi.org/10.1113/jphysiol.2014.272518

17 Perlman, I. Kinetics of bleaching and regeneration of rhodopsin in abnormal (RCS) and normal albino rats in vivo. *The Journal of Physiology* **278**, 141-159 (1978). https://doi.org:https://doi.org/10.1113/jphysiol.1978.sp012297

18 Zhang, P., Goswami, M., Zawadzki, R. J. & Pugh, E. N., Jr. The Photosensitivity of Rhodopsin Bleaching and Light-Induced Increases of Fundus Reflectance in Mice Measured In Vivo With Scanning Laser Ophthalmoscopy. *Invest. Ophthalmol. Vis. Sci.* **57**, 3650-3664 (2016). https://doi.org:10.1167/iovs.16-19393

19 Lamb, T. D. Photoreceptor spectral sensitivities: Common shape in the long-wavelength region. *Vision Res.* **35**, 3083-3091 (1995). https://doi.org:https://doi.org/10.1016/0042-6989(95)00114-F

20 Morgan, J. I. W. & Pugh, E. N., Jr. Scanning Laser Ophthalmoscope Measurement of Local Fundus Reflectance and Autofluorescence Changes Arising from Rhodopsin Bleaching and Regeneration. *Invest. Ophthalmol. Vis. Sci.* **54**, 2048-2059 (2013). https://doi.org:10.1167/iovs.12-11089

21 Pandiyan, V. P., Nguyen, P. T., Pugh, E. N. & Sabesan, R. Human cone elongation responses can be explained by photoactivated cone opsin and membrane swelling and osmotic response to phosphate produced by RGS9-catalyzed GTPase. *Proceedings of the National Academy of Sciences* **119**, e2202485119 (2022). https://doi.org:doi:10.1073/pnas.2202485119

22 Norren, D. V. & Vos, J. J. Spectral transmission of the human ocular media. *Vision Res.* **14**, 1237-1244 (1974). https://doi.org:https://doi.org/10.1016/0042-6989(74)90222-3

23 Mahroo, O. A. R. & Lamb, T. D. Recovery of the human photopic electroretinogram after bleaching exposures: estimation of pigment regeneration kinetics. *The Journal of Physiology* **554**, 417-437 (2004). https://doi.org:https://doi.org/10.1113/jphysiol.2003.051250

24 WA, H. The quantum efficiency of bleaching of rhodopsin in situ. *The Journal of Physiology* **129**, 22-23P (1955).

25 Dowling, J. E. & Hubbard, R. Effect of Instantaneous Flashes on Adaptation of the Eye: Effects of Brilliant Flashes on Light and Dark Adaptation. *Nature* **199**, 972-975 (1963). https://doi.org:10.1038/199972a0

26 Williams , T. P. Photoreversal of Rhodopsin Bleaching. *J. Gen. Physiol.* **47**, 679-689 (1964). https://doi.org:10.1085/jgp.47.4.679

27 Hofmann, K. P. & Lamb, T. D. Rhodopsin, light-sensor of vision. *Prog. Retin. Eye Res.* **93**, 101116 (2023). https://doi.org:https://doi.org/10.1016/j.preteyeres.2022.101116
